# Chimeric antigen receptor T cell therapy targeting CD94 for cytotoxic T/NK cell malignancies

**DOI:** 10.64898/2026.11.12.765663

**Authors:** Cao Cuong Le, Jinsheng Weng, Jingwei Liu, Federico Mario Aletti, Xiaoyun Cheng, Yongfu Tang, Sridevi Patchva, Matthew Richard, Maya Balakumaran, Mario Marques-Piubelli, Luisa Solis, Wei Lu, Chun-Yen Lin, Fuliang Chu, Jingjing Cao, Abdur Rehman, Kamini Singh, Owhofasa Agbedia, Sushanth Gouni, Karan Chohan, Faezeh Darbaniyan, Selin Kolsuz, Kumudha Balakrishnan, Neeraj Saini, Michael R. Green, Christopher R Flowers, Swaminathan P. Iyer, Francisco Vega, Sattva S Neelapu

## Abstract

Chimeric antigen receptor (CAR) T cell therapy against pan-B-cell antigens is highly effective, but analogous approaches for T-cell lymphomas risk prolonged T-cell aplasia. We developed a CAR-T therapy targeting a post-thymic antigen, CD94, a receptor with restricted expression in normal tissues and immune cells but enriched in cytotoxic T/NK-cell malignancies. Second generation CD94 CAR T cells generated using a specific and high-affinity anti-CD94 monoclonal antibody showed potent and selective cytotoxicity against CD94-positive tumor cells *in vitro* and completely eradicated NK-cell leukemia xenografts and patient-derived xenograft models of hepatosplenic T-cell lymphoma and NK/T-cell lymphoma *in vivo*. Incorporation of truncated B-cell maturation antigen (BCMA) as a safety switch enabled rapid elimination of CAR T cells using anti-BCMA T-cell engagers or anti-BCMA antibody-drug conjugate. Super-resolution microscopy demonstrated intracellular sequestration of CD94 by the CAR, preventing fratricide. Collectively, these findings support anti-CD94 CAR-T therapy as a promising and novel treatment for cytotoxic T/NK-cell neoplasms.

## INTRODUCTION

Chimeric antigen receptor (CAR) T-cell therapy targeting the pan-B-cell antigen, CD19, has revolutionized the treatment of relapsed/refractory B-cell lymphomas and leukemias.^1,2^ However, development of CAR-T therapy for T/NK-cell malignancies, which represent 10-15% of all lymphoid neoplasms has been challenging.^3^ These malignancies are characterized by highly aggressive clinical behavior with long-term survival observed in less than 20% of patients.^4,5^ Current treatment options, including combination chemotherapy and hematopoietic stem cell transplantation, often provide limited benefit due to substantial toxicity and inadequate efficacy, underscoring the need for novel, safer, and more effective therapeutic approaches.

CAR-T therapies targeting pan-T-cell antigens such as CD5 and CD7 have been explored for T-cell malignancies. However, unlike B-cell-directed CAR-T therapies, these approaches carry the risk of profound and prolonged T-cell aplasia because the targeted antigens are broadly expressed on normal T cells.^6–9^ Consequently, patients may develop severe immunodeficiency and life-threatening infections, which unlike B-cell deficiency, could lead to fatal infections unless rescued by allogeneic stem cell transplantation.^10,11^ Additionally, shared antigen expression between CAR T cells and malignant T cells can result in fratricide, limiting CAR T cell expansion, persistence, and overall therapeutic efficacy. Therefore, targeting novel antigens that have a restricted expression profile amongst normal T cells is critical for developing effective CAR-T therapies for T-cell malignancies while minimizing fratricide and preserving protective immune function.^12^

T/NK-cell malignancies comprise a heterogeneous group of diseases that can be classified according to their cell of origin, reflecting the stage of T-cell development within the thymus or peripheral immune system and their mature functional subsets.^13^ The cell of origin largely determines the biological, genetic, and clinical features of the major T/NK-cell malignancy subtypes. We therefore hypothesized that CAR-T therapies targeting post-thymic antigens expressed on the cells of origin of these malignancies could provide a safe and effective therapeutic strategy. Here, we describe the development of a novel second-generation CAR-T product targeting CD94 (KLRD1), a C-type lectin receptor expressed on 50-70% of mature NK cells and γδ T cells, as well as a subset of cytotoxic CD8⁺ T cells amongst normal lymphocytes.^14,15^ CD94 functions as either an activating or inhibitory receptor through heterodimerization with NKG2C/E or NKG2A, respectively.^16,17^ Previous studies have demonstrated high CD94 expression across multiple cytotoxic T/NK-cell neoplasms, including extranodal NK/T-cell lymphoma, hepatosplenic T-cell lymphoma, primary cutaneous γδ T-cell lymphoma, aggressive NK-cell leukemia, primary cutaneous CD8⁺ aggressive epidermotropic cytotoxic T-cell lymphoma, subcutaneous panniculitis-like T-cell lymphoma, monomorphic epitheliotropic intestinal T-cell lymphoma, T-cell large granular lymphocytic leukemia (T-LGL), and chronic lymphoproliferative disorder of NK cells (CLPD-NK, also known as NK-LGL).^18–22^ We therefore developed a novel CD94-targeted CAR T cell therapy and evaluated its safety and antitumor efficacy in preclinical models.

## RESULTS

### CD94 expression in normal tissues and T/NK-cell malignancies

To assess the safety of targeting CD94, we examined its expression in normal human tissues by immunohistochemistry (IHC). Evaluation of 32 normal tissues demonstrated CD94 expression in occasional lymphocytes within secondary lymphoid organs, such as the tonsils, with little to no expression detected in most non-hematopoietic tissues, including the brain, heart, lung, liver, kidney, pancreas, and intestines (**Extended Data Fig. 1a**). Flow cytometric analysis of peripheral blood mononuclear cells (PBMCs) from five healthy donors confirmed CD94 expression on 50– 70% of NK cells and γδ T cells, and on fewer than 15% of CD8⁺ T cells, while expression was largely absent on CD4⁺ T cells, B cells, monocytes, and dendritic cells (**Extended Data Fig. 1b-f**). The median CD94 antigen density was approximately 20-fold lower on normal CD8⁺ T cells than on NK leukemia cell lines. In contrast, consistent with prior reports,^18–22^ CD94 was highly and uniformly expressed in most NK/T-cell lymphomas, hepatosplenic T-cell lymphomas, and a subset of other T-cell lymphoma subtypes (**Extended Data Fig. 1g-h**). Collectively, the restricted expression of CD94 in normal immune cells and its high expression in cytotoxic T/NK-cell malignancies support its potential as a safe target for CAR T cell therapy.

### Characterization of an anti-human CD94 monoclonal antibody and generation of anti-CD94 CAR T cells

To identify a suitable binder for CD94-directed CAR T cells, we generated an anti-human CD94 mouse monoclonal antibody using standard hybridoma technology (**Extended Data Fig. 2a-b**). This antibody specifically bound CD94-expressing NK leukemia cell lines (NKL and KHYG-1), but not their CD94-knockout counterparts. Consistent with the known expression profile of CD94, the antibody bound a large fraction of NK cells and a smaller subset of T cells from healthy donor PBMC, while showing minimal binding to B cells or monocytes, indicating high specificity (**Fig. 1a**). To further confirm specificity, we screened the antibody against a human membrane proteome array comprising approximately 6,000 proteins. The antibody bound exclusively to KLRD1 (CD94), either as a monomer or in heterodimeric complexes with its canonical binding partners KLRC1 (NKG2A), KLRC2 (NKG2C), and KLRC3 (NKG2E), with no detectable binding to other targets (**Fig. 1b**). Bio-layer interferometry demonstrated high-affinity binding, with an equilibrium dissociation constant (K_D_) of approximately 1 nM (data not shown).

**Figure 1.**
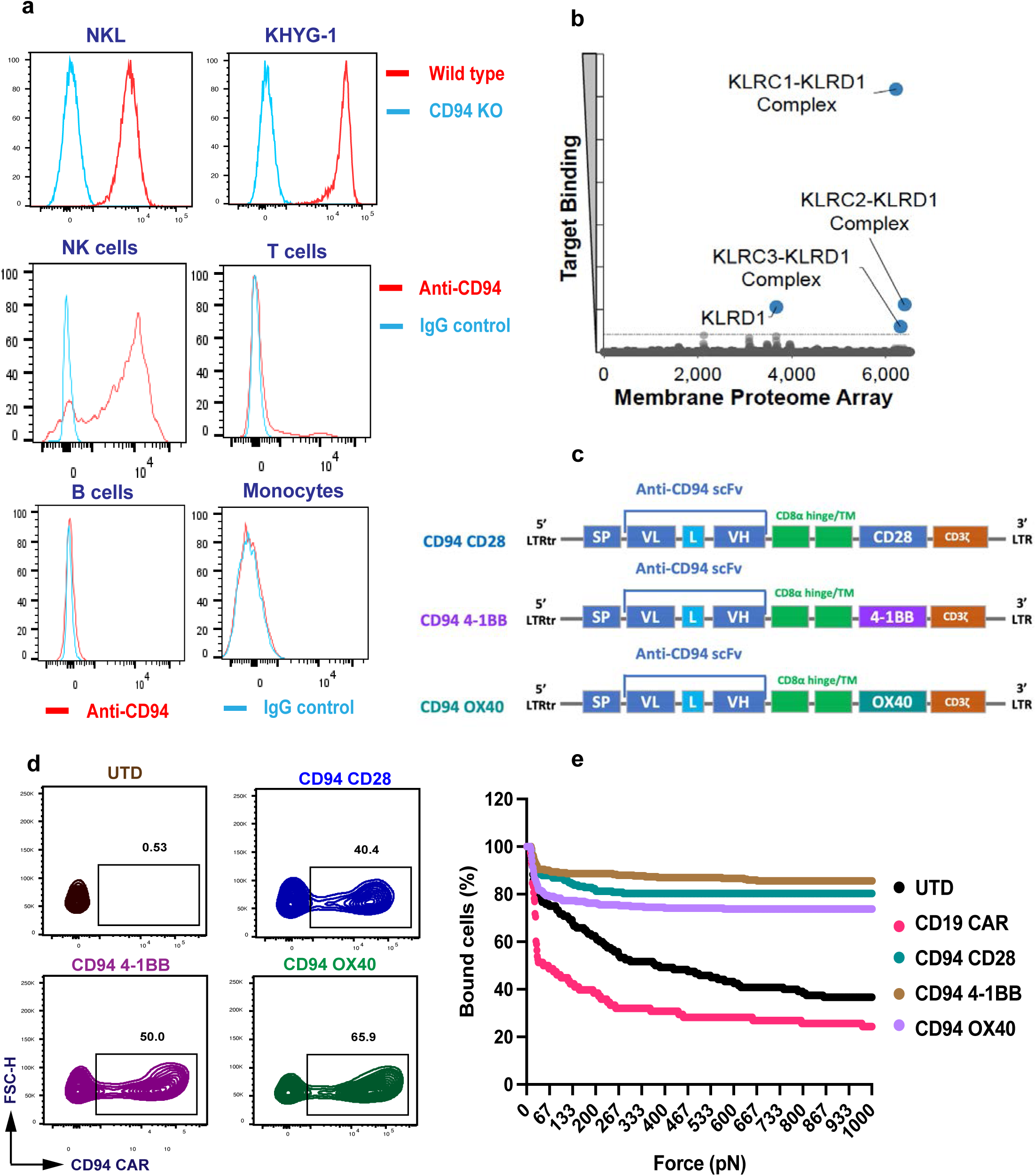
Anti-CD94 antibody and CAR generation. **a)** Binding of anti-CD94 monoclonal antibody generated by us to wild-type NKL and KHYG-1 NK-cell leukemia cell lines and their isogenic CD94 knock-out (KO) counterparts and human peripheral blood NK cells, T cells, B cells, and monocytes was assessed by flow cytometry. An IgG isotype antibody was used as a control. **b)** Off-target binding of the anti-CD94 antibody was assessed by flow cytometry against ∼6,000 human membrane proteins expressed in HEK-293T cells in duplicates. The Y-axis represents the ‘Target Binding’ calculated for each target in the membrane proteome array (MPA) library. The X-axis represents the indexed locations of each of the proteins in the MPA Library; showing the numerical order in which each target or well is read (which includes duplicates, controls, and empty wells). Each dot represents one unique protein clone from the MPA library. Blue dots represent targets that were bound. The dotted line represents 3 standard deviations above the background. **c)** Design of three CD94-targeting CAR constructs. **d)** CD94-targeting CAR transduction efficiency in primary T cells was assessed by flow cytometry by staining with recombinant human CD94 protein. **e)** Avidity of various CD94-targeting CAR T cells to KHYG-1 target cells was assessed by z-Movi. CD19 CAR-T and UTD T cells were used as controls.

We next generated three second-generation anti-CD94 CAR constructs incorporating a CD94-specific single-chain variable fragment (scFv) derived from the monoclonal antibody, a CD8α hinge and transmembrane domain, and a CD3ζ signaling domain, combined with either a CD28, 4-1BB, or OX40 costimulatory domain (**Fig. 1c**). Following lentiviral transduction of primary human T cells, robust surface expression of all CAR constructs was confirmed by demonstrating binding to recombinant human CD94 protein (**Fig. 1d**). Avidity analyses showed strong binding of all three anti-CD94 CAR-T products to CD94-positive KHYG-1 target cells, with no significant differences observed among the constructs (**Fig. 1e**), indicating comparable target engagement.

### Antitumor activity of CD94 CAR T cells *in vitro*

To evaluate the antitumor activity *in vitro*, we co-cultured the anti-CD94 CAR T cells with CD94-positive NKL and KHYG-1 target cells. All three CAR-T constructs induced potent cytotoxicity, eliminating tumor cells with high efficiency across both tumor cell lines over two days (**Fig. 2a-b**). Repeated challenge with tumor cells over two weeks showed that all three CAR-T products were capable of serial cytotoxicity (**Fig. 2c-d**). Furthermore, the CAR T cells also efficiently eliminated KHYG-1 tumor clones with up to 19-fold lower CD94 antigen density than the wild-type cell line but had no significant cytotoxicity against CD94 knock-out tumor cells, demonstrating antigen-dependent tumor recognition and killing (**Fig. 2e-f** and **Extended Data Fig. 3a-b**).

**Figure 2.**
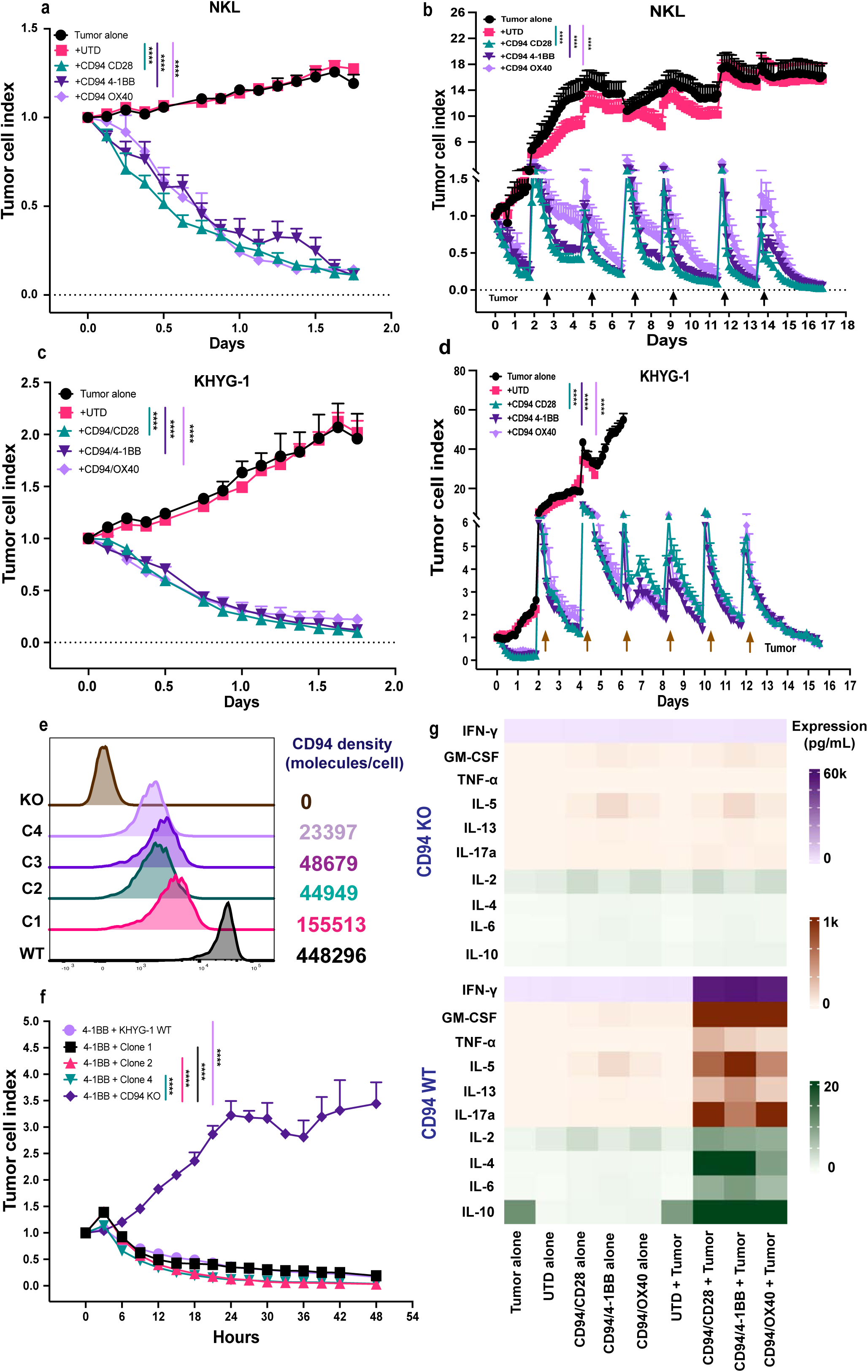
Cytotoxic activity and cytokine expression by CD94-targeting CAR T cells *in vitro*. **a-d)** Cytotoxic activity by CD94-targeting CAR T cells against NKL (**a** and **b**) and KHYG-1 (c and d) NK-cell leukemia cell lines at an Effector: Target (E:T) ratio of 1:2 was monitored by serial imaging using the Incucyte Live-Cell Analysis System over two days (a and c) or over 16-17 days in a serial cytotoxicity assay (**b** and **d**). **e)** KHYG-1 clones with varying CD94 expression and antigen density or CD94 knock-out. **f)** Cytotoxic activity of CD94-targeting CAR T cells (4-1BBz) against KHYG-1 clones with varying CD94 antigen density. **g)** Cytokine expression by CD94-targeting CAR T cells after 48 hours in response to KHYG-1 wild type (WT) or CD94^KO^ KHYG-1 cells is shown as a heatmap. Tumor alone and UTD T cells are shown as controls where applicable. Statistical significance was determined using one-way ANOVA with Dunnett’s multiple comparisons test. ****: p < 0.0001. All experiments were performed in triplicate wells, and the data represent mean ± standard error of mean where applicable. Results shown are representative of at least five separate experiments using CAR T cells generated from five different donors.

Cytokine release assays further showed that all three CD94 CAR T cell products secreted high levels of interferon-γ and other effector cytokines when co-cultured with CD94-positive target cells, whereas cytokine production remained at background levels in the absence of target antigen (**Fig. 2g** and **Extended Data Fig. 3c**). Collectively, these findings demonstrate that CD94 CAR T cells mediate robust and selective antitumor activity against CD94-expressing malignancies, including tumors with relatively low antigen density, supporting their potential therapeutic application in CD94+ neoplasms.

### Antitumor activity of CD94 CAR T cells *in vivo*

To evaluate the *in vivo* efficacy of CD94 CAR T cells, NSG mice were engrafted with either NKL lymphoma cells or KHYG-1 NK leukemia cells. One week later, mice received a single intravenous dose of CD94 CAR T cells or untransduced T cells and were monitored longitudinally by bioluminescence imaging (**Fig. 3a**). CD94 CAR T cell treatment resulted in rapid and sustained tumor clearance in both xenograft models whereas progressive tumor growth was observed in control-treated and untreated animals (**Fig. 3b-c** and **e-f**). To assess therapeutic activity against tumors expressing lower levels of CD94, we also evaluated efficacy of CD94 CAR T cells in a KHYG-1 CD94-low model. Despite reduced antigen expression, CD94 CAR T cells retained potent antitumor activity, resulting in complete tumor eradication in treated mice compared with control groups (**Extended Data Fig. 4a-b**). Consistent with the *in vitro* findings, all three CAR constructs demonstrated comparable antitumor efficacy, with no significant differences in tumor clearance or overall survival among the CAR-T treatment groups across all three tumor models. No significant weight loss or other external signs of toxicity were observed in any CAR-T-treated cohort (**Fig. 3d** and **g** and **Extended Data Fig. 4c**).

**Figure 3.**
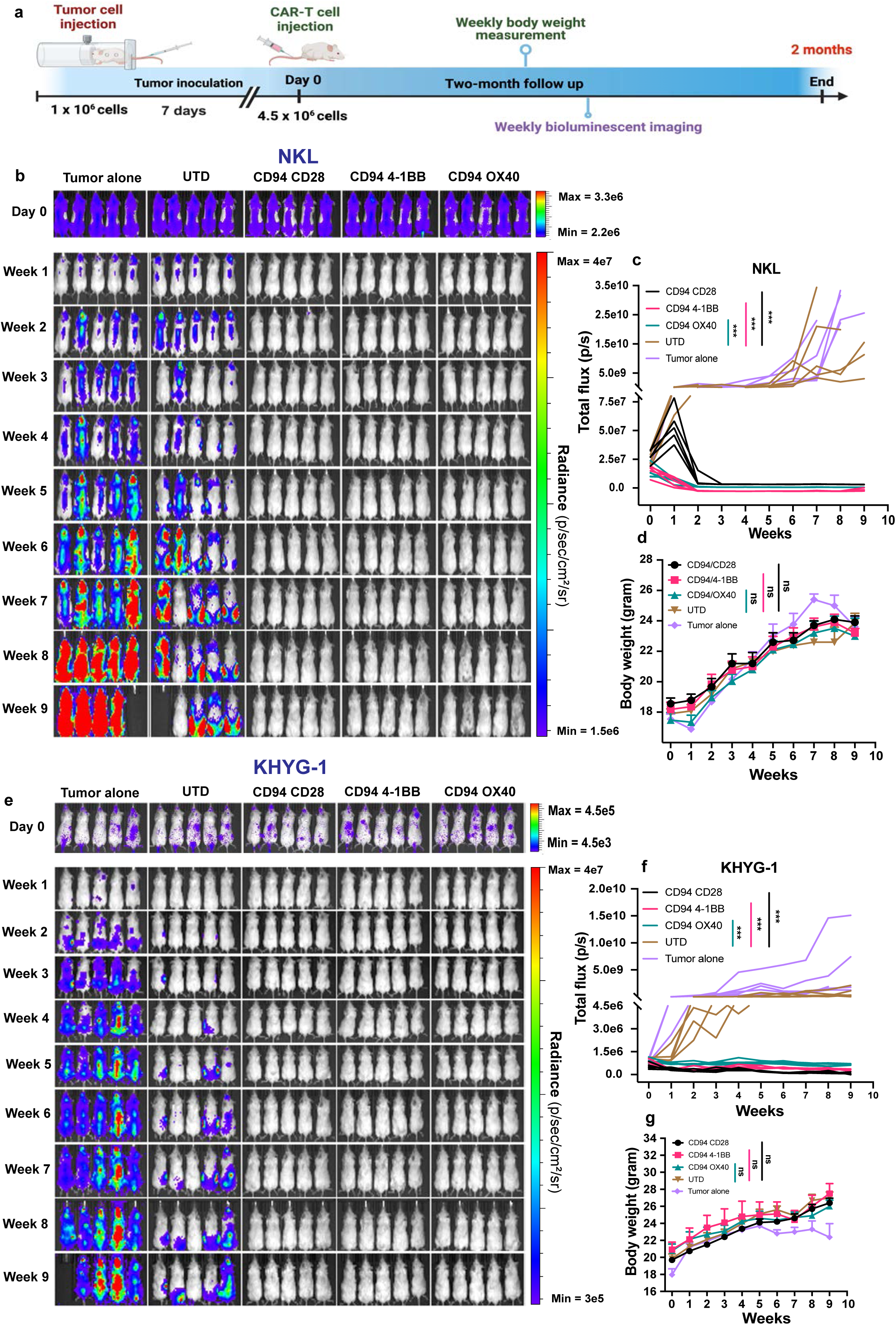
Antitumor efficacy of CD94-targeting CAR T cells *in vivo*. **a)** Experimental outline to assess efficacy of CD94 CAR T cells without a safety switch against NKL and KHYG-1 cell line-derived xenograft (CDX) models in NSG mice. **b-d)** Bioluminescence imaging (**b**), total flux (**c**), and body weight (**d**) in NKL CDX model. **e-g)** Bioluminescence imaging (**e**), total flux (**f**), and body weight (g) in KHYG-1 CDX model. Total flux values were calculated from normalized radiance spectra (p/s/cm²/sr) using Living Image software. Data are represented individually for each mouse. Statistical significance was determined using one-way ANOVA with Dunnett’s multiple comparisons test (**: p < 0.005). Results shown are representative of at least three separate experiments for each CDX model using CAR T cells generated from three different donors. UTD, untransduced; p/s, photons/second; ns, not significant.

Because no significant differences in antitumor efficacy were observed among the three CD94 CAR constructs (CD28, 4-1BB, and OX40), we selected the 4-1BB-containing CAR for further development based on its favorable safety profile and clinical track record in FDA-approved CAR T cell therapies.^23^ To enhance safety, we incorporated a truncated B-cell maturation antigen (tBCMA) molecule into the CD94-4-1BB CAR construct separated by self-cleaving peptide (**Fig. 4a**). Following lentiviral transduction with this bicistronic construct, dual expression of both the CAR and tBCMA was observed on primary T cells, with the CAR surface expression levels comparable to those observed with the CD94 CAR lacking the safety switch (**Fig. 4c**). Antitumor efficacy in a KHYG-1 cell line xenograft model was comparable between CAR T cells with or without the safety switch confirming that incorporation of the tBCMA safety switch did not impair the function of CAR T cells (**Fig. 4b and 4d-e**).

**Figure 4.**
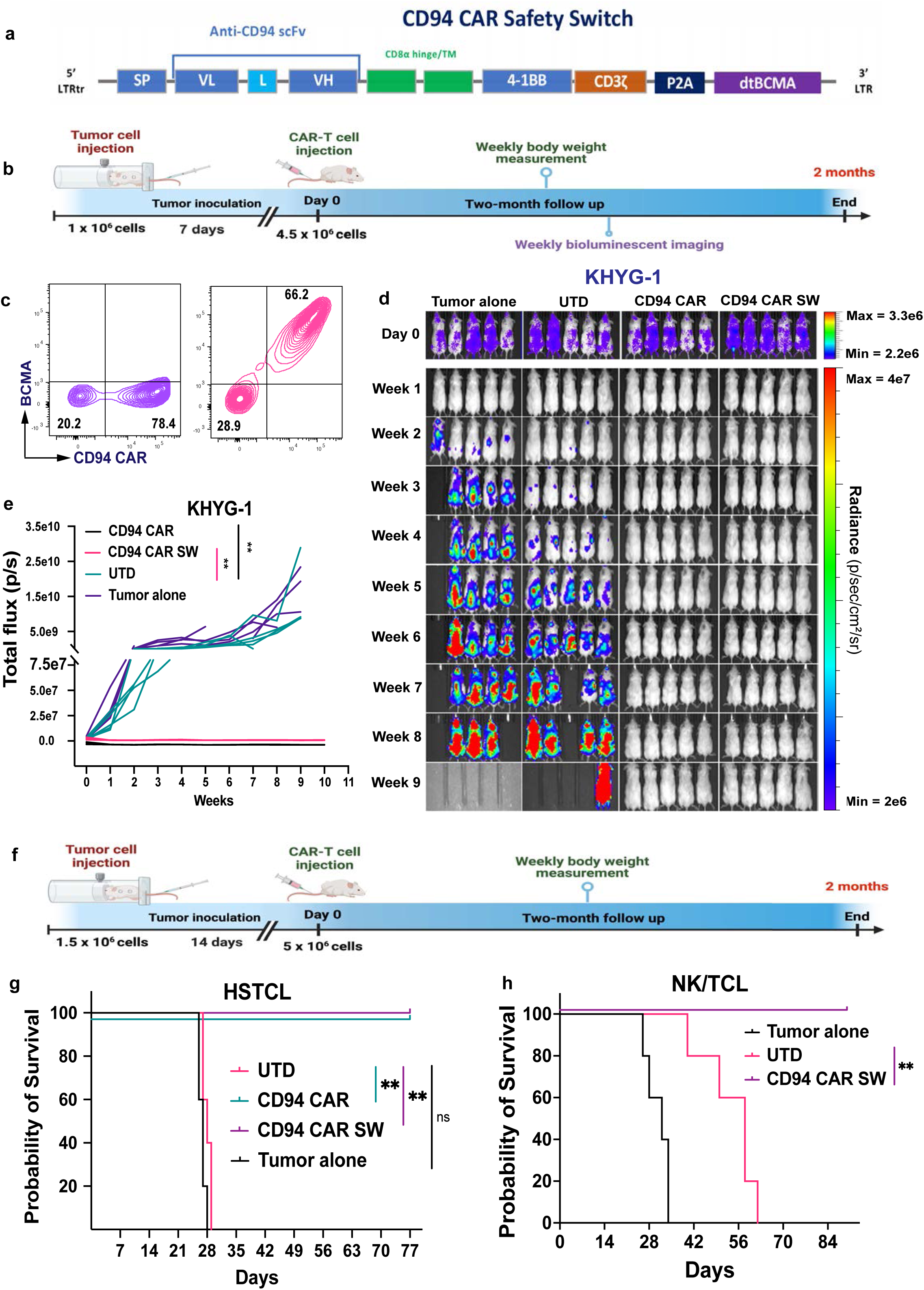
Antitumor efficacy of CD94-targeting CAR T cells with dtBCMA safety switch *in vivo*. **a)** Design of CD94 CAR construct with or without dtBCMA safety switch. **b)** Experimental outline to assess efficacy of CD94 CAR T cells with dtBCMA safety switch against KHYG-1 CDX models in NSG mice. **c)** Transduction efficiency in primary T cells transduced with CD94 CAR without (*left*) or with dtBCMA safety switch (*right*). Expression of CD94 CAR and dtBCMA safety switch is shown. **d** and **e)** Bioluminescence imaging (**d**) and total flux (**e**) in KHYG-1 CDX model. Total flux values were calculated from normalized radiance spectra (p/s/cm²/sr) using Living Image software. Data are represented individually for each mouse. **f)** Experimental outline to assess efficacy of CD94 CAR T cells with or without dtBCMA safety switch against patient-derived xenograft (PDX) models of hepatosplenic T cell lymphoma (HSTCL) and NK/T-cell lymphoma (NK/TCL) in NSG mice. **g** and **h)** Survival of mice in HSTCL (**g**) and NK/TCL (**h**) PDX models. Statistical significance for total flux was determined using one-way ANOVA with Dunnett’s multiple comparisons test (**: p < 0.005). Statistical analysis in PDX models was determined using log-rank (Mantel-Cox) test (**: p < 0.005). Results shown are representative of at least three separate experiments for both CDX and PDX models using CAR T cells generated from three different donors. dtBCMA, dual truncated BCMA; CDX, cell line-derived xenograft; UTD, untransduced; SW, safety switch; p/s, photons/second; ns, not significant.

To further evaluate therapeutic efficacy, we tested CD94 CAR T cells in two patient-derived xenograft (PDX) models, a hepatosplenic T-cell lymphoma and an NK/T-cell lymphoma. In both tumor models, CD94 CAR T cells significantly prolonged survival compared to the control-treated or untreated animals (**Fig. 4f-h** and **Extended Data Fig. 5a-b**). Importantly, no evidence of graft-versus-host disease or weight loss was observed in any cohort (**Extended Data Fig. 5c**). Evaluation of mice in the hepatosplenic T-cell lymphoma model at study termination showed marked splenomegaly in tumor alone and untransduced T cell groups as compared with the CAR-T-treated groups (**Extended Data Fig. 5d**). Consistent with this, flow cytometric analysis confirmed presence of tumor cells in the liver, spleen, and bone marrow of control mice but no tumor cells were detected in the mice treated with CD94 CAR T cells (**Extended Data Fig. 5e**). Together, these results demonstrate the potent antitumor activity of CD94 CAR T cells against CD94+ malignancies, including tumors with low antigen density and in clinically relevant PDX models of cytotoxic T/NK-cell malignancies.

### Rapid manufacturing enhances CD94 CAR T cell efficacy

Recent studies indicate that rapid manufacturing of CAR T cells preserves a less-differentiated, naïve and stem-cell memory phenotype and yields a product with greater anti-tumor potency, while significantly reducing cost of goods, logistical hurdles, and vein-to-vein time.^24^ Here, we compared the phenotype and antitumor activity of CD94 CAR T cells generated over 3 days versus those generated over 10 days (**Fig. 5a**). Phenotypically, Day 3 CAR T cells retained a higher proportion of naïve-like T cells compared with Day 10 CAR T cells (**Fig. 5b**). To determine whether these phenotypic differences translated into improved antitumor activity, Day 3 and Day 10 CD94 CAR T cells were compared in serial tumor rechallenge assays. Although both products efficiently eliminated tumor cells after multiple rounds of tumor challenges at 1:2 effector to target (E:T) ratio, only Day 3 CAR T cells maintained sustained cytotoxic activity at 1:5 E:T ratio consistent with improved potency (**Fig. 5c-d**). The enhanced activity of rapidly manufactured CD94 CAR T cells was further validated *in vivo*. In NSG mice bearing NKL tumor xenografts, Day 3 CAR T cells induced durable tumor regression and improved survival compared with Day 10 CAR T cells at low doses of 0.2 to 0.5 × 10^6^ CAR^+^ T cells/mouse without any signs of weight loss or toxicity (**Fig. 5e-i** and **Extended Data Fig. 6a-c**). Collectively, these findings demonstrate that rapid manufacturing preserves a less differentiated T-cell state and enhances the antitumor potency of CD94 CAR T cells, supporting the clinical development of an accelerated manufacturing approach.

**Figure 5.**
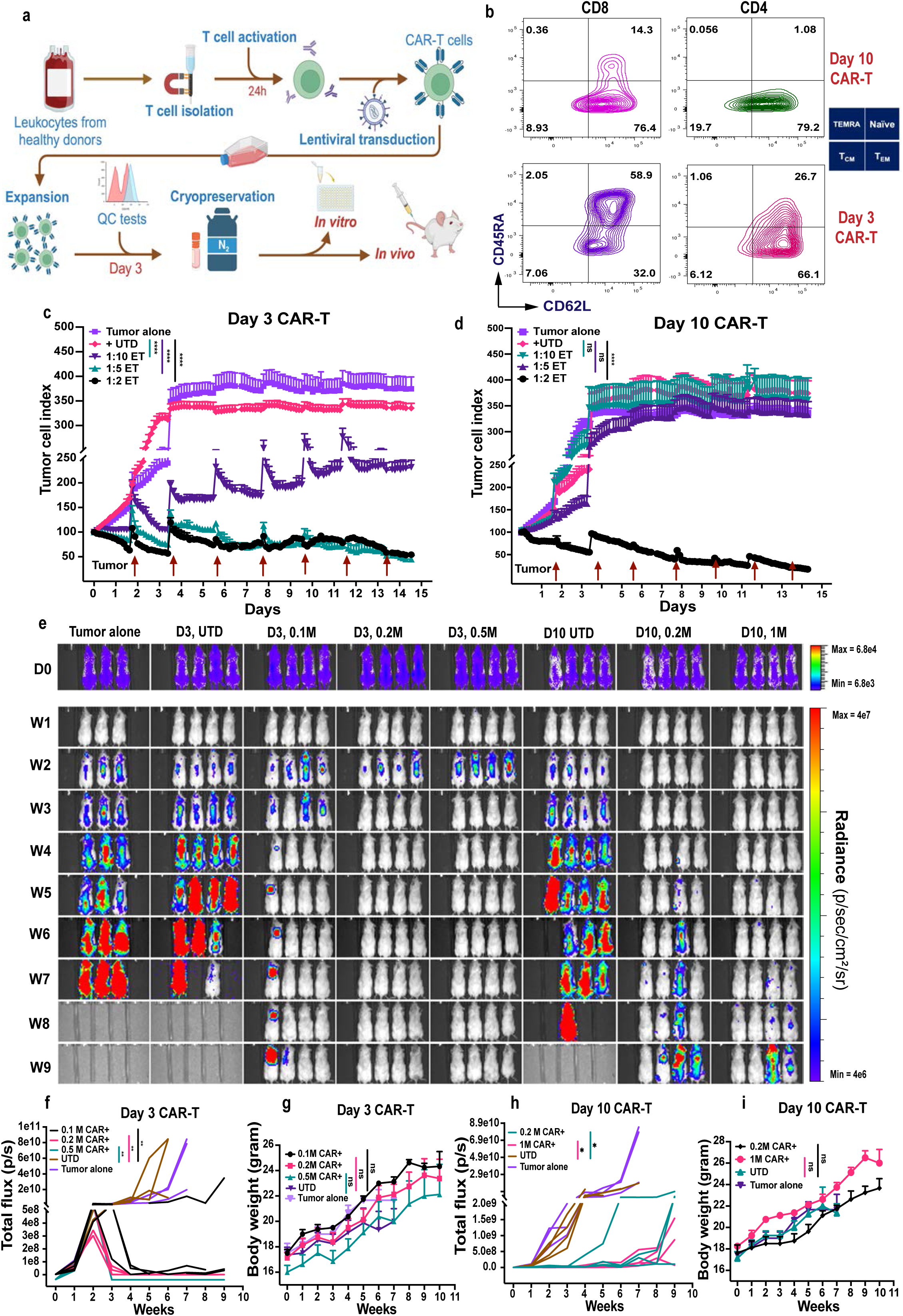
Antitumor activity of rapidly manufactured CD94-targeting CAR T cells. **a)** Schematic illustrating the workflow for rapidly manufactured CD94 CAR T cells. **b)** Phenotype of CD4 and CD8 CAR T cells manufactured over 3 days vs. 10 days. **c** and **d)** Serial cytotoxic activity against NKL of CD94-targeting CAR T cells manufactured over 3 days (**c**) vs. 10 days (**d**) at 1:2, 1:5, and 1:10 E:T ratios. The experiment was performed in triplicate wells and data are represented as mean ± standard error of mean. Results shown are representative of at least two separate experiments. Statistical significance was determined using one-way ANOVA with Dunnett’s multiple comparisons test (****: p < 0.0001, ns: not significant). **e-i)** NKL cells were injected intravenously via the tail vein into NSG mice at a dose of 1.5 × 10⁶ cells per mouse and allowed to engraft for seven days. NKL tumor-bearing mice were then treated with Day 3 CAR T cells at 0.1, 0.2, and 0.5 × 10⁶ cells per mouse or with Day 10 CAR T cells at 0.2 and 1 × 10⁶ cells per mouse. Day 3 and day 10 untransduced T cells were injected into each mouse in the control groups at 0.5 × 10⁶ and 1 × 10⁶ cells, respectively. Bioluminescence imaging (**e**), total flux (**f** and **h**), and body weight (**g** and **i**) are shown for the different groups. Total flux values were calculated from normalized radiance spectra (p/s/cm²/sr) using Living Image software. Data are represented for each individual mouse. Statistical significance was determined using one-way ANOVA with Dunnett’s multiple comparisons test (**: p < 0.005). Results shown are representative of at least three separate experiments.

### Assessment of on-target, off-tumor cytotoxicity by CD94 CAR T cells

Given the restricted expression of CD94 on subsets of normal NK cells and T cells, we evaluated the potential for on-target, off-tumor toxicity of CD94 CAR T cells toward healthy immune cell populations. Co-culture of autologous CD94 CAR T cells with healthy donor PBMC showed partial elimination of NK cells as compared to control PBMC alone or PBMC co-cultured with untransduced T cells (**Fig. 6a-b**). This was further confirmed by lower fold-expansion of GFP-transduced NK cells cultured in the presence of IL-2 and CD94 CAR T cells as compared to control groups (**Fig. 6c**). To assess the impact on normal T-cell populations, purified total T cells were cultured alone or with autologous CD94 CAR T cells. Consistent with the expression of profile of CD94, we observed susceptibility of CD94^+^ subsets to CD94 CAR T cell-mediated killing in TCRαβ, CD8^+^, and γδ T cells, but negligible cytotoxicity against CD4^+^ T cells (**Fig. 6d-g**). Importantly, despite this selective depletion, the absolute number of T cells recovered by day 6 in each of these subsets and was comparable to control T cell group except for γδ T cells, which showed only partial recovery (**Fig. 6h-k**).

**Figure 6.**
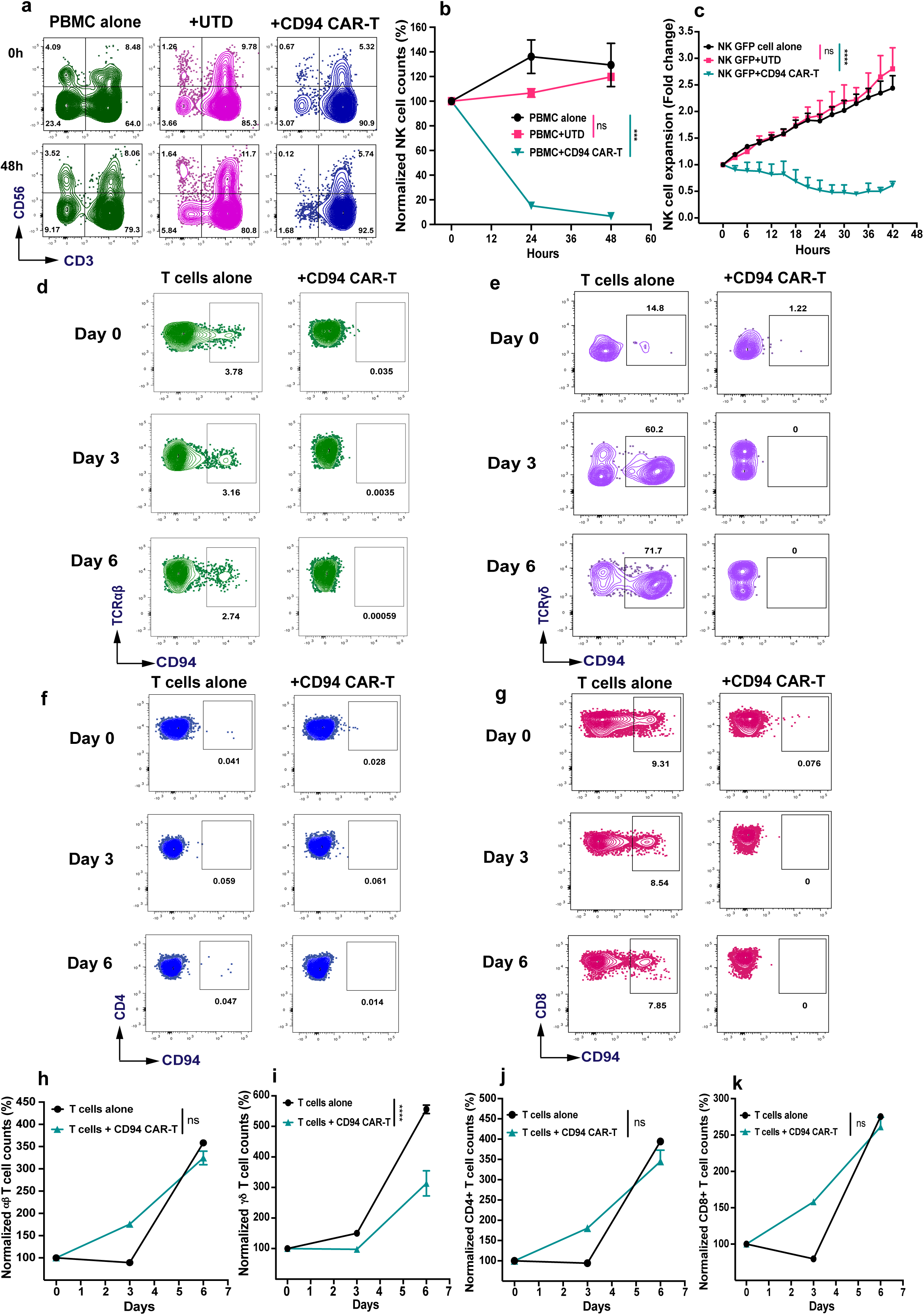
Effects of CD94-targeting CAR T cells against immune cell subsets. **a** and **b)** CD94 CAR T cells and activated untransduced (UTD) T cells were co-cultured with autologous PBMCs for two days and effects on NK cell numbers were determined by flow cytometry at baseline, 24 hours, and 48 hours using absolute counting beads. Representative data (**a**) and data from triplicate well are shown (**b**). Statistical significance was determined by one-way ANOVA with Dunnett’s multiple comparisons test (***: p < 0.001, ns: not significant). **c)** Healthy donor-derived NK cells were transduced with GFP and co-cultured with activated UTD or CD94 CAR T cells. NK-GFP cell viability and expansion were monitored using Incucyte Live-Cell Analysis System for up to 42 hours. Statistical significance was determined by one-way ANOVA with Dunnett’s multiple comparisons test (****: p < 0.0001, ns: not significant). **d-g)** Activated T cells were cultured with or without CD94 CAR T cells and the effects on CD94⁺ *a*β T cells (**d**), CD94⁺ *γδ* T cells (**e**), CD94⁺ CD4 T cells (**f**), and CD94⁺ CD8 T cells (**g**) were determined by flow cytometry and absolute counting beads at days 0, 3, and 6. **h-k)** Normalized cell counts (%) were calculated from absolute cell counts for total *a*β T cells (**h**), total *γδ* T cells (**i**), total CD4 T cells (**j**), and total CD8 T cells (**k**). Statistical analysis was performed for the day 6 time point for all groups using Student’s t-test (****: p < 0.0001, ns: not significant).

To further confirm the on-target, off-tumor effects, we assessed the peripheral blood immune compartment in CD34⁺ hematopoietic stem cell-engrafted NSG-hu-IL-15 humanized mice following treatment with CD94 CAR T cells. As compared to control untreated group, CD94 CAR-T treatment resulted in elimination of most of the NK cells and a small subset of T cells that were CD94^+^ (**Extended Data Fig. 7a-c**). In agreement with this, the T cell compartment was largely preserved compared to baseline whereas significant NK cell depletion was observed in CD94 CAR-T-treated group (**Extended Data Fig. 7d-g**). Collectively, these data indicate that CD94 CAR T cells induce predictable and selective depletion of CD94-expressing lymphocyte populations particularly NK cells and γδ T cells while preserving most of the adaptive immune compartment.

### Efficacy of tBCMA safety switch

To evaluate the functionality of the tBCMA safety switch, the tBCMA^+^ CD94 CAR T cells were treated *in vitro* with anti-BCMA bispecific T-cell engaging antibodies, teclistamab or elranatamab, or with an anti-BCMA antibody-drug conjugate, belantamab mafodotin. Treatment with any one of these agents resulted in rapid and significant depletion of tBCMA^+^ CD94 CAR T cells whereas untreated cells, isotype-control-treated cells, and/or CD94 CAR T cells lacking tBCMA were unaffected (**Fig. 7a-e**, **Extended Data Fig. 8a-b, and Extended Data Fig. 9a**). Importantly, these effects were observed with each of these agents at concentrations expected to be achieved in the serum with standard dosing in patients.

**Figure 7.**
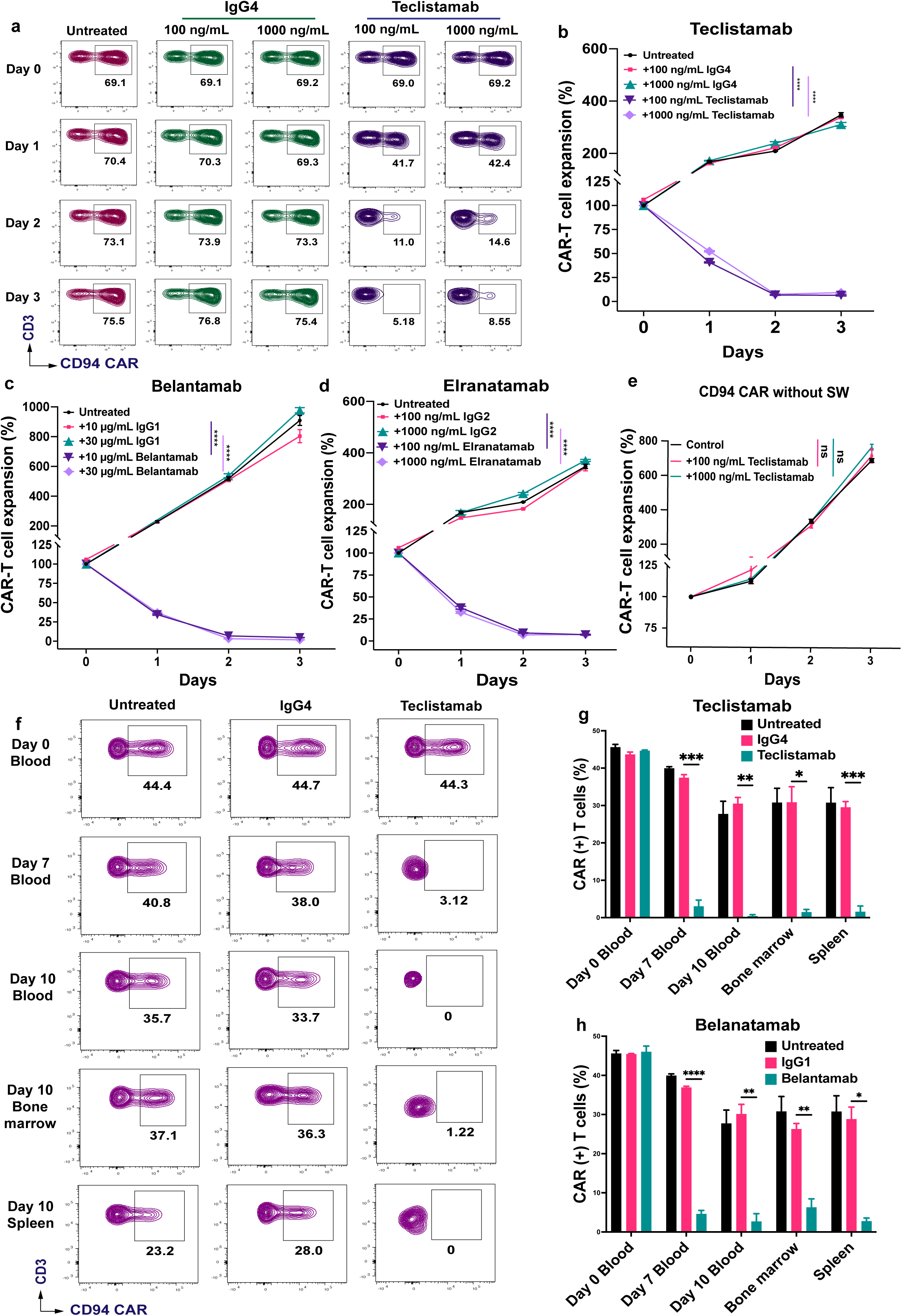
Efficacy of dtBCMA safety switch. **a-e)** CD94 CAR T cells harboring a dtBCMA safety switch construct were untreated or treated at the indicated concentrations with teclistamab (**a-b**), belantamab mafodotin (**c**), or elranatamab (**d**) for three days or their respective isotype IgG antibody controls. The number of CAR T cells at each indicated time point was determined by flow cytometry and absolute counting beads. Percent change in absolute number of CAR T cells compared to baseline is shown over 3 days. **e)** Effects of teclistamab on CD94 CAR T cells lacking the dtBCMA safety switch. Statistical significance was determined using one-way ANOVA with Dunnett’s multiple-comparisons test (****, p < 0.0001; ns, not significant). All experiments were performed in triplicate wells and the data represent mean ± standard error of mean. Results shown are representative of at least three separate experiments using CAR T cells generated from three different donors. **f-h)** CD94 CAR T cells with dtBCMA safety switch were intravenously injected into NSG mice via the tail vein at a dose of 4 × 10⁶ cells per mouse. Seven day later, mice were untreated or treated with teclistamab or its IgG4 isotype control antibody at a dose of 1.5 mg/kg (f and g) or belantamab mafodotin or its IgG1 isotype control at a dose of 2.5 mg/kg (h). CAR T cell percentage was determined by flow cytometry serially at the indicated time points in peripheral blood or in bone marrow and spleen at necropsy. Statistical analysis was performed using two-way ANOVA with Dunnett’s multiple-comparisons test (****, p < 0.0001; ***, p < 0.001; **, p < 0.01; *, p < 0.05; ns, not significant).

To assess the efficacy of the tBCMA safety switch *in vivo*, tBCMA^+^ CD94 CAR T cells were intravenously infused into non-tumor-bearing NSG mice and three days later, mice were treated with a single dose of teclistamab, belantamab mafodotin, or their respective isotype control antibodies. Peripheral blood, spleen, and bone marrow samples were collected at serial time points to quantify CAR T cell persistence. Flow cytometric analysis seven days later demonstrated marked depletion of tBCMA^+^ CD94 CAR T cells following treatment with either BCMA-targeting agent compared with isotype-treated or untreated controls (**Fig. 7f** and **Extended Data Fig. 9b**). The depletion was sustained in the blood, bone marrow, and spleen when the mice were assessed at necropsy on day 10 (**Fig. 7g-h**). Together, these results establish that clinically available BCMA-directed therapies can serve as an effective pharmacologic safety switch, enabling rapid and sustained elimination of tBCMA^+^ CD94 CAR T cells in case of unexpected toxicities.

### Resistance to fratricide by CD94 CAR T cells

Since expression of CD94 on CAR T cells may lead to fratricide, we determined whether CD94 is upregulated on activated CAR T cells. Repeated stimulation of CD19 CAR T cells with a CD19^+^ B-cell lymphoma cell line every 3 days resulted in progressive upregulation of CD94 in CD19 CAR T cells and an increase in CD94+ CAR T cell number with each successive round of tumor stimulation (**Fig. 8a-b**). This is consistent with the notion that CD94 is expressed together with NKG2A on cytotoxic T cells and the heterodimer functions as an immune checkpoint on activated CD8^+^ T cells.^16,17^ However, stimulation of CD94-targeting CAR T cells repeatedly with CD94^+^ KHYG-1 tumor cells did not similarly result in CD94 upregulation (**Fig. 8a**).

**Figure 8.**
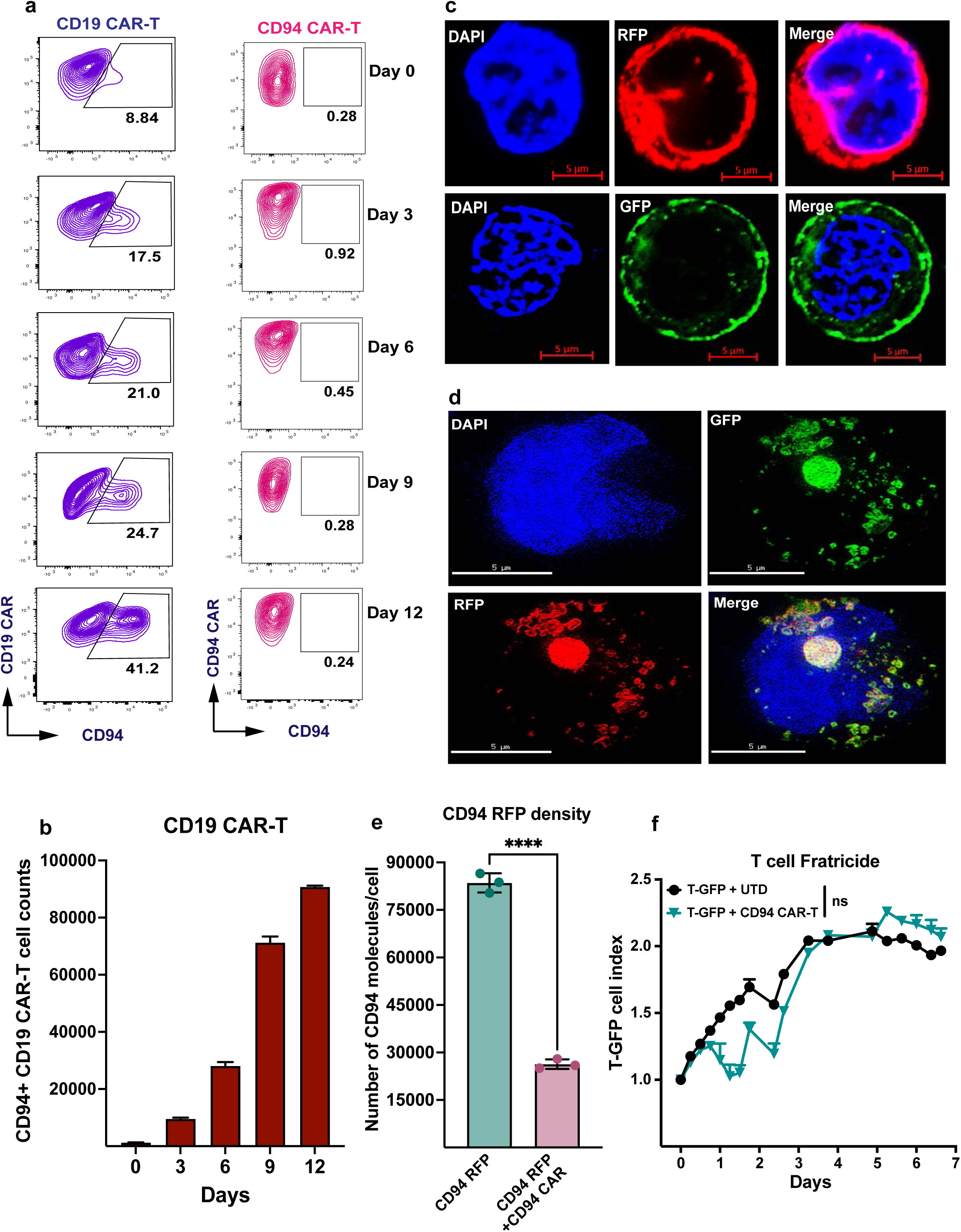
Evaluation of CD94 CAR T cell fratricide. **a)** CD19-targeting and CD94-targeting CAR T cells were co-cultured with Jeko-1 and KHYG-1 tumor cells, respectively, at 1:1 E:T ratio, and target tumor cells were added to each co-culture every three days for up to 12 days. CD94 expression on the CAR T cells was determined by flow cytometry prior to each tumor rechallenge. **b)** Graph showing change in absolute numbers of CD94⁺ CD19 CAR T cells over the 12 days. **c)** K562 cells were transduced with CD94-RFP (upper panels) and CD94 CAR-GFP (lower panels) to express fluorescent fusion proteins and confocal microscopy imaging was performed using Airyscan with fixed cells. **d)** K562 cells transduced with CD94-RFP were sorted, and then transduced with CD94 CAR-GFP to co-express both fusion fluorescent proteins. Imaging was performed using super-resolution Structured Illumination Microscopy (SIM) with fixed cells. **e)** Membrane CD94 density in K562 cells co-expressing CD94-RFP and CD94 CAR-GFP was measured using flow cytometry with standardized quantum microsphere beads. **f)** Activated T cells lentivirally transduced with GFP were sorted based on high GFP expression and co-cultured with CD94-targeting CAR T cells or untransduced (UTD) T cells from the same donor. The GFP-transduced T cell (T-GFP) index was measured using Incucyte Live-Cell Analysis System. Statistical significance was determined for days 5 to 7 using two-way ANOVA with Dunnett’s multiple comparisons test. ns: not significant.

To elucidate the mechanism underlying this phenomenon, we lentivirally transduced CD94-RFP or CD94-targeting CAR-GFP fusion proteins into K562 cells and found that they localized to the cell membrane as expected using structured illumination microscopy (**Fig. 8c**). However, introduction of CD94-targeting CAR-GFP into K562 cells expressing CD94-RFP resulted in loss of surface localization of CD94 and extensive intracellular colocalization of CD94 and the CAR, suggesting sequestration of CD94 by the CAR within intracellular compartments (**Fig. 8d**). Quantification of CD94 surface expression before and after CAR transduction by flow cytometry demonstrated marked reduction in surface CD94 expression in co-transduced K562 cells as compared to K562 transduced with CD94-RFP alone (**Fig. 8e**). Thus, the CAR-mediated sequestration likely explains the absence of CD94 upregulation in stimulated CD94 CAR T cells and suggests an intrinsic safeguard against fratricide.

Because low-level CD94 expression persists on a subset of normal CD8⁺ T cells, we next evaluated whether CD94 CAR T cells could eliminate normal T cells over time. GFP-labeled primary human T cells were co-cultured with autologous untransduced or CD94 CAR T cells and monitored longitudinally by live-cell imaging. Although transient CD8⁺ T cell depletion was observed during the initial phase of co-culture, residual CD8⁺ T cells persisted and subsequently expanded, resulting in stable cell numbers comparable to control cultures at later time points (**Fig. 8f**). These findings suggest that only a minor subset of normal CD8⁺ T cells is susceptible to CD94 CAR T cell-mediated killing.

Collectively, these results demonstrate that intracellular sequestration of CD94 by the CAR effectively prevents fratricide, enabling sustained expansion and function of CD94 CAR T cells. Furthermore, the limited impact on normal T cell populations supports a favorable safety profile and highlights the therapeutic potential of CD94-targeted CAR-T therapy for cytotoxic T/NK-cell malignancies.

## DISCUSSION

Our findings demonstrate that CD94 is a promising target for CAR T-cell therapy in CD94-expressing T/NK-cell leukemias and lymphomas. We confirmed the specificity of our proprietary anti-CD94 antibody through comprehensive screening against the human membrane proteome and validation using isogenic cell lines. In addition, cytotoxicity mediated by CD94 CAR T cells was shown to be strictly antigen dependent. Across multiple cell line-derived and patient-derived xenograft models, CD94 CAR T cells induced robust antitumor activity, resulting in complete eradication of widely disseminated disease. Consistent with previous reports,^24^ CAR T cells manufactured using a 3-day process exhibited superior potency compared with products generated over 10 days. Collectively, these data support the potential clinical application of CD94 CAR T-cell therapy in patients with relapsed/refractory CD94-expressing T/NK-cell neoplasms including extranodal NK/T-cell lymphoma, hepatosplenic T-cell lymphoma, primary cutaneous γδ T-cell lymphoma, aggressive NK-cell leukemia, primary cutaneous CD8⁺ aggressive epidermotropic cytotoxic T-cell lymphoma, subcutaneous panniculitis-like T-cell lymphoma, and monomorphic epitheliotropic intestinal T-cell lymphoma among others.^18–22^ Although individually rare, these disorders collectively represent a patient population with a substantial unmet medical need. Notably, patients with hepatosplenic T-cell lymphoma or aggressive NK-cell leukemia are typically diagnosed at a young age, with median survival ranging from only 6 to 18 months despite currently available therapies.^25,26^ Likewise, outcomes remain poor for patients with advanced or refractory extranodal NK/T-cell lymphoma.^27^ Thus, an effective CD94-directed CAR T-cell therapy has the potential to significantly improve outcomes for these otherwise highly lethal malignancies.

The development of CAR T-cell therapies for T-cell malignancies has been particularly challenging. Most strategies have focused on broadly expressed T-cell antigens, such as CD5 and CD7, analogous to targeting pan-B-cell antigens including CD19, CD20, and CD79b in B-cell malignancies.^1,28–32^ While such approaches have broad applicability, they carry the inherent risk of prolonged pan-T-cell aplasia and the associated morbidity and mortality from severe infections. Furthermore, because these target antigens are expressed on the therapeutic T cells themselves, manufacturing generally requires genetic disruption of the target antigen to prevent fratricide, thereby increasing manufacturing complexity, cost, and potential genotoxicity.

To overcome these limitations, we selected CD94, a post-thymic differentiation antigen with restricted expression on normal immune cells but frequent expression in malignancies derived from cytotoxic lymphocytes, including NK cells, γδ T cells, and CD8⁺ T cells. This expression pattern provides an opportunity to selectively target malignant cells while minimizing on-target, off-tumor toxicity. Consistent with this premise, our *in vitro* studies using human peripheral blood mononuclear cells and our *in vivo* studies in humanized mice demonstrated depletion of only CD94-expressing subsets of NK cells, γδ T cells, and a small fraction of CD8⁺ T cells. Importantly, CD94-negative populations within each compartment were spared, thereby avoiding the pan-T-cell aplasia associated with many alternative targets. Although CD94 expression can be induced on activated CAR T cells, our data demonstrate that fratricide is effectively prevented through sequestration of CD94 by the CAR itself. Together, these findings suggest that targeting post-thymic, lineage-restricted antigens associated with the cell of origin may represent a safer and more selective strategy for CAR T-cell therapy in T/NK-cell malignancies.

The degree of immune-cell depletion anticipated with CD94-directed therapy is likely to be clinically manageable. Patients with primary immunodeficiencies characterized predominantly by NK-cell deficiency are principally susceptible to recurrent herpesvirus infections, which can generally be managed through vaccination, antiviral prophylaxis, and targeted antiviral therapies.^33,34^ To further enhance safety, we incorporated a novel dtBCMA safety switch into the CAR construct. Unlike established suicide-switch systems, including inducible caspase-9,^35^ HSV-TK,^36^ and EGFRt,^37^ specialized activating agents, potentially immunogenic transgenes, or less predictable antibody-mediated depletion mechanisms, the dtBCMA platform can be activated using multiple FDA-approved anti-BCMA therapeutics already available in clinical practice. In our studies, a single administration of either teclistamab or belantamab mafodotin induced rapid and sustained depletion of CD94 CAR T cells from both peripheral blood and tissues, demonstrating an efficient mechanism for eliminating engineered cells in the event of severe toxicity. Importantly, recent clinical data further support the safety of targeting CD94. In a phase 1 study, DR-01, a non-fucosylated human IgG antibody directed against CD94 that mediates target-cell depletion primarily through antibody-dependent cellular cytotoxicity, was well tolerated in patients with cytotoxic T/NK-cell lymphomas.^38^ Taken together, the restricted expression profile of CD94, the limited impact on normal immune compartments, and the incorporation of the dtBCMA safety switch support a favorable therapeutic index for this approach.

Several limitations should be acknowledged. First, CD94-directed CAR T-cell therapy is applicable only to malignancies that express CD94 and therefore will not address the full spectrum of T-cell neoplasms. Second, in patients with a leukemic phase of disease, contamination of the leukapheresis product by malignant cells may result in inadvertent CAR transduction, potentially leading to antigen masking and tumor escape.^39^ Addressing this challenge may require development of an allogeneic CD94 CAR T-cell platform. Third, activation of the dtBCMA safety switch for management of severe cytokine release syndrome or immune effector cell-associated neurotoxicity syndrome could preclude subsequent use of anti-BCMA T-cell engagers because of overlapping toxicities. In such circumstances, however, belantamab mafodotin may remain a viable option for CAR T cell depletion. Despite these limitations, the profound unmet clinical need and poor outcomes associated with cytotoxic T/NK-cell malignancies strongly support continued development of this therapeutic strategy.

In conclusion, this study establishes CD94 as a selective and clinically actionable target for CAR T-cell therapy in cytotoxic T/NK-cell malignancies. CD94 CAR T cells demonstrated potent and durable antitumor activity across multiple *in vitro* and *in vivo* models, including tumors with relatively low antigen density, while largely sparing normal immune-cell populations. Fratricide was minimal due to intrinsic sequestration of CD94 by the CAR construct, and inclusion of the dtBCMA safety switch provides an additional safeguard to enhance clinical safety. Collectively, these findings provide a strong preclinical rationale for clinical translation and support the ongoing phase 1 evaluation of an autologous anti-CD94 CAR T-cell product in patients with relapsed or refractory CD94-expressing T/NK-cell neoplasms (NCT07382817).

## ACKNOWLEDGEMENTS

This study was supported by the University of Texas MD Anderson Cancer Center T/NK-cell Malignancies Moonshot (SSN and FV), Department of Defense grant (CA220435 to SSN), Cancer Prevention Research Institute of Texas Multi-Investigator Research Award (RP240439 to SSN), and National Institutes of Health NCI Cancer Center Support Grant to the University of Texas MD Anderson Cancer Center (P30 CA016672) (Flow Cytometry and Cellular Imaging Facility). The MD Anderson Lymphoma Tissue Bank is supported by KW Cares Foundation.

## AUTHORSHIP CONTRIBUTIONS

CCL performed the CAR subcloning, designed and/or performed the *in vitro* and *in vivo* experiments, analyzed the data, prepared the figures, and wrote the manuscript; JW generated and screened hybridomas, helped with antibody and CAR subcloning, designed the *in vitro* and *in vivo* experiments; JL, YT, and SP generated the lentiviral plasmid, developed the BCMA safety switch and designed the *in vitro* and *in vivo* experiments; FA, XC, MR, MB, CYL, FC, JC, SK, AR, KS, OA, SG, KC, SK, KB, NS, MRG, CRF, and SI assisted with the design of *in vitro* and *in vivo* experiments and provided critical feedback; MMP, LS, WL, and FV performed IHC staining and analyzed the data of the patient biospecimens; FD supervised and performed data analysis; SSN conceived the study, designed the *in vitro* and *in vivo* experiments, analyzed data, obtained funding and wrote the manuscript. All authors reviewed, provided critical feedback, and approved the final manuscript.

