## Supplemental File for "Chimeric antigen receptor T cell therapy targeting CD94 for cytotoxic T/NK cell malignancies"

#### ONLINE METHODS

##### Ethics statement

All healthy donor and patient samples used in this study were deidentified. All experiments using human samples were approved by the Institutional Review Board at The University of Texas MD Anderson Cancer Center and conducted in accordance with the Declaration of Helsinki. Mice were housed and maintained at 25 °C under a 12-hour light-dark cycle. All animal studies were conducted in compliance with the guidelines of the Institutional Animal Care and Use Committee (IACUC) of The University of Texas MD Anderson Cancer Center under an approved protocol.

##### Blood and tumor samples

Healthy donor PBMCs were isolated from buffy coats obtained from Gulf Coast Regional Blood Center. Formalin fixed paraffin-embedded sections were generated from archived tumor tissue blocks obtained from T/NK cell lymphoma patients. Histological tissue slides of normal human tissues were purchased from TissueArray.Com LLC while NK/T-cell lymphoma tissue slides were purchased from Quickarrays, Inc. All human samples used in this study were deidentified and approved by the Institutional Review Board at The University of Texas MD Anderson Cancer Center.

##### Cell lines and cell culture

KHYG-1 cells were purchased from the German Collection of Microorganisms and Cell Cultures GmbH (DSMZ). NKL cells were provided by Dr. Jerome Ritz (Harvard Medical School). NK cell lines were cultured in complete RPMI-1640 medium (Gibco) supplemented with 10% FBS (R&D Systems), 100 units/mL Penicillin-Streptomycin (Gibco), and 10 ng/mL IL-2. K562 and 293T cells, obtained from ATCC, were cultured in IMDM and DMEM (Gibco), respectively, each supplemented with 10% FBS and 100 units/mL Penicillin-Streptomycin (Gibco). Primary T cells and CAR T cells were cultured in RPMI-1640 supplemented with 300 IU/mL IL-2, 10% FBS (R&D Systems), 100 units/mL Penicillin-Streptomycin (Gibco), 25 mM HEPES buffer (Gibco), 1 mM Sodium Pyruvate (Gibco), 1% MEM Non-Essential Amino Acids Solution (100X) (Gibco), 100 µg/mL Normocin (InvivoGen), 0.1% β-mercaptoethanol (v/v) (Gibco). All human primary cells and cell lines were maintained at 37°C in a humidified incubator with 5% $CO_2$. Patient-derived xenograft (PDX) of hepatosplenic T-cell lymphoma was obtained from The Public Repository of Xenograft (PRoXe). These PDX-derived cells were maintained and expanded through *in vivo* propagation in NSG mice.

#### Cell line transduction

KHYG-1 and NKL wild-type cells were co-transduced with two lentiviral vectors encoding luciferase-GFP and IL-15-RFP to generate stable cell lines expressing both luciferase and IL-15, intended for *in vitro* and *in vivo* experiments. KHYG-1 IL-15 and NKL IL-15 cells were cultured in complete RPMI-1640 medium without cytokine supplementation. K562 cells were transduced with lentiviral vectors encoding either CD94-RFP or CD94 CAR-GFP fusion proteins to establish single fusion protein-expressing lines. Subsequently, CD94-RFP-expressing K562 cells were further transduced with a lentiviral vector encoding CD94 CAR-GFP, resulting in a stable K562 cell line co-expressing both CD94-RFP and CD94 CAR-GFP fusion proteins. GFP and RFP fluorescence were used as reporters for single cell clone selection.

#### Flow cytometry

Cells were blocked with human Fc receptor (FcR) blockers for 15 minutes at room temperature and then stained with appropriate fluorochrome-conjugated antibodies and live/dead dyes, followed by flow cytometry analysis using the BD LSRFortessa. Flow cytometry data were analyzed using FlowJo version 10 and/or R/RStudio.

#### Antigen density quantification

Cells were stained with antibodies of interest following standard blocking and washing procedures. Standardized quantum microsphere beads (Bangs Laboratories) were stained with each individual antibody in separate tubes according to the manufacturer's instructions. Both stained cells and beads were analyzed using the BD LSRFortessa, applying the same instrument settings for consistency. Flow cytometry data were analyzed using FlowJo version 10. Antigen density was calculated based on geometric mean fluorescence intensity using the QuickCal® tool (Bangs Laboratories) provided by the manufacturer.

#### Monoclonal antibody generation

Human CD94 cDNA (GenBank: U30610.1) was cloned into a lentiviral vector and expressed in a murine fibroblast cell line (L cells). BALB/c mice (Taconic Biosciences) were immunized with CD94-expressing L cells following protocols described in a previous study [1], and a CD94-specific monoclonal antibody was subsequently generated using standard hybridoma technology at the Monoclonal Antibody Core Facility of MD Anderson Cancer Center.

#### Membrane Protein Array

The anti-CD94 antibody was tested for binding specificity using a membrane protein array containing 6,000 surface proteins (Integral Molecular).[2]

**CAR constructs**

Second-generation chimeric antigen receptor (CAR) constructs targeting CD94 were synthesized by incorporating the CD94 scFv derived from the novel murine anti-CD94 monoclonal antibody with CD8α hinge and transmembrane domains, one of three distinct co-stimulatory domains CD28, 4-1BB, or OX40, and a CD3ζ signaling domain. Each of the three CD94 CAR constructs was cloned into a lentiviral vector backbone under the control of the MSCV promoter. A safety switch previously developed by us, dual truncated B-cell maturation antigen (dtBCMA)[3,4], was incorporated into the CD94/4-1BB CAR construct, separated by a P2A skip element encoding a self-cleaving peptide.

**Lentivirus generation**

Lentivirus encoding CD94 CAR constructs was produced in 293T cells by co-transfecting CD94 CAR transfer plasmids with the ABM 3rd Generation Packaging System Mix using Lipofectamine 3000 (Invitrogen). Lentiviral supernatants were collected at 24 and 48 hours post-transfection and filtered through a 0.45 µm cellulose acetate filter. Based on lentiviral titers, the filtered supernatant was either snap-frozen directly or concentrated using Lenti-X Concentrator (Takara). Concentrated lentivirus was resuspended in Opti-MEM medium (Gibco) and stored at −80 °C for long-term use.

**CAR T cell Generation**

Peripheral blood mononuclear cells (PBMCs) were isolated using Histopaque-1077 (Sigma) and cryopreserved at −80 °C in CS10 cryopreservation medium (STEMCELL Technologies). T cells were subsequently isolated from thawed PBMCs using the Pan T Cell Isolation Kit (Miltenyi Biotec). Purified T cells were activated using the ImmunoCult™ Human CD3/CD28/CD2 T Cell Activator (STEMCELL Technologies) for 24 hours in complete RPMI-1640 medium as described previously. Lentiviral transduction was then performed in 24-well plates using either 15 µg/well of Retronectin (Takara) for the viral prebound method, or 10 µg/mL of Vectofusin-1 (Miltenyi Biotec) for the viral unbound method. Activated T cells were seeded at $1 \times 10^6$ cells/well and spin-transduced with CD94 CAR lentivirus at 1000 × g, 32 °C for 1 hour (prebound method) or 2 hours (unbound method). Transduction efficiency was evaluated on day 2 post-transduction by flow cytometry. Afterward, CD94 CAR T cells were expanded in complete RPMI-1640 medium supplemented with 300 IU/mL recombinant human IL-2 and additional reagents. Cells were split every 2 days to maintain a density of 1 × $10^6$ cells/mL. CAR T cells were harvested and cryopreserved either on day 2 post-transduction (for shortened manufacturing protocols) or on day 7 (for conventional manufacturing protocols). Donor-matched untransduced (UTD) T cells, served as controls, were activated and cultured under the same conditions as the CD94 CAR T cells.

**Cell avidity assay**

Target (KHYG-1) and effector (CAR T/UTD) cells were cultured and maintained to achieve cell viability greater than 90%. Prior to the cell avidity assay, both cell types were incubated overnight in fresh complete medium. Target cells were then seeded onto the flow cell of the z-Movi Cell Avidity Analyzer (Lumicks) to form a cell monolayer with at least 90% confluency. Cells were incubated at 37 °C for 30 minutes to promote firm adherence to the glass surface of the flow cell, and the medium was exchanged every hour thereafter. Effector cells were labeled using CellTrace™ Far Red dye (Invitrogen) according to the manufacturer's instructions. Stained effector cells were then resuspended at a density of 10 × $10^6$ cells/mL in complete RPMI-1640 medium with 10% FBS and gently loaded onto the prepared monolayer of target cells for binding, with the number of effector cells limited to 200 - 500 per run. Effector cells were allowed to bind to the target cells for 5 minutes before the avidity measurement. Following incubation, an acoustic force gradient was applied using the z-Movi Cell Avidity Analyzer ranging from 0 to 1000 pN. The avidity between effector and target cells was measured based on the percentage of bound effector cells remaining out of the total loaded cells, corresponding to the applied acoustic force.

**Incucyte Imaging**

All co-culture experiments using the Incucyte Live-Cell Imaging System were performed in 96-well plates pre-coated with either soluble Poly-D-Lysine (Gibco) or Poly-L-Ornithine (Sigma), following the manufacturers' instructions. GFP- and/or RFP-expressing target cells were co-cultured with unlabeled CD94 CAR T cells or donor-matched UTD T cells at various effector-to-target (E:T) ratios. For serail cytotoxicity assay, the culture medium was exchanged every other day, and fresh medium containing new viable tumor cells was added to each well for up to two weeks. Scanning was performed at the indicated time points to monitor changes in tumor cell signal density.

**Cytokine release assay**

KHYG-1 wild-type (WT) and CD94 knockout ($CD94^{KO}$) KHYG-1 cells were co-cultured with CD94 CAR T cells or UTD T cells at a 1:1 E:T ratio for 24 hours at 37°C with 5% $CO_2$. Monocultures of CD94 CAR T and UTD T cells were included as controls. Following incubation, the supernatant was collected and cryopreserved at −80°C until further analysis. Multiplex enzyme-linked immunosorbent assay (multiplex ELISA) assays were performed in triplicate using customized cytokine array plates (Meso Scale Diagnostics), each well containing captured spots for 10 cytokines including IL-2, IL-4, IL-5, IL-6, IL-10, IL-13, IL-17A, GM-CSF, TNF-α, and IFN-γ. All procedures were carried out according to the manufacturer's instructions.

##### Confocal microscope imaging

K562 cells expressing either CD94-RFP, CD94 CAR-GFP, or co-expressing both fusion proteins were seeded onto coverslips (Warner Instruments) with a thickness of 0.17 ± 0.02 mm and incubated at 37°C in a 5% $CO_2$ incubator for 6 hours. The cells were then fixed and stained with DAPI for nuclear visualization. Confocal microscopy using Airyscan was performed with a Zeiss LSM 880 confocal microscope (Zeiss) to detect CD94-RFP and CD94 CAR-GFP fusion proteins separately. Co-localization of CD94-RFP and CD94 CAR-GFP was analyzed using Structured Illumination Microscopy (SIM) on the OMX Blaze V4 super-resolution microscope.

##### Immunohistochemistry

Immunohistochemistry (IHC) staining was performed using the Leica BOND™ RX autostainer and the BOND™ Polymer Refine Detection Kit (Leica Biosciences, Cat# DS9800). A human tissue array (Tissuearray.com, Cat# FDA999w3) containing 32 different organs from three healthy donors was stained with a commercial anti-CD94 antibody (Abcam, clone EPR21003, Cat# ab235441) at 1:400 dilution. All incubation steps were carried out at room temperature with wash cycles between steps unless otherwise specified. Antibody-antigen binding was visualized using a brown DAB Refine chromogen for 10 minutes, followed by counterstaining with hematoxylin for 8 minutes. Slides were subsequently dehydrated, cleared in xylene, and cover-slipped using Cytoseal™ XYL mounting medium.

##### Mouse studies

Female NOD.Cg-$Prkd^{cscid}$ $Il2rg^{tm1Wjl}$/SzJ (NSG) mice, aged 4-6 weeks, were used to establish cell line-derived xenografts (CDX) with luciferase-expressing NKL and KHYG-1 cells, as well as patient-derived xenograft (PDX) of hepatosplenic T-cell lymphoma. Female NOD.Cg-$Prkdc^{scid}$ $Il2rg^{tm1Wjl}$ Tg(IL15)1Sz/SzJ mice engrafted with human $CD34^+$ hematopoietic cord blood stem cells (humanized mice) were used to evaluate the safety of CD94 CAR T cells by assessing potential off-target cytotoxicity against human immune cells. All mice were obtained from the Jackson Laboratory. Animal care, tumor burden, and graft-versus-host disease monitoring were performed daily by the laboratory staff and/or the Department of Veterinary Medicine and Surgery (DVMS) at the University of Texas MD Anderson Cancer Center. Euthanasia procedures were conducted in accordance with DVMS recommendations to ensure compliance with ethical animal care standards.

**Bioluminescence imaging**

Mice bearing luciferase-expressing tumor cells were anesthetized using isoflurane in accordance with the DVMS-approved standard operating protocol. Following anesthesia, mice were intraperitoneally injected with 150 mg/kg of D-Luciferin (Promega, USA). Bioluminescence imaging was performed using the IVIS Spectrum System (Perkin Elmer) at the indicated time points.

**Data analysis and statistics**

Confocal microscopy data were processed using ZEN Blue version 3.10 (Zeiss), while IHC images were processed using Aperio ImageScope version 12 (Leica Biosystems). Bioluminescent signal quantification was performed using Living Image version 4.7. Graphs and statistical visualizations were generated using GraphPad Prism version 10. Area under the curve (AUC) values were calculated for experiments involving multiple timepoints, followed by statistical analyses. Survival analysis was conducted using the Kaplan-Meier method, with comparisons using the log-rank (Mantel-Cox) test. Other statistical analyses were performed using Student's t-test, one-way ANOVA, or two-way ANOVA with Dunnett's multiple comparisons test, depending on the dataset and experimental designs.

#### EXETENDED FIGURE LEGENDS

**Extended Data Figure 1. CD94 expression in normal and tumor tissues. a)** Immunohistochemical staining of normal human tissues using a commercial anti-CD94 antibody. Normal tonsil and NK/T lymphoma served as positive controls. **b)** Uniform Manifold Approximation and Projection (UMAP) showing cell clusters of immune cell subsets from five healthy donor PBMC (left panel) and CD94 expression in various clusters (right panel). **c)** CD94 expression in immune cell subsets (n = 5) was quantified by flow cytometry with standardized quantum microsphere beads and visualized as heatmap for surface CD94 density (molecules/cell) and as bubble plot for percentage of cells with CD94 expression. NKL and KHYG-1 NK-cell leukemia cells served as positive controls. **d-f)** Percent cells with CD94 expression (**d** and **f**) and CD94 density (molecules/cell) (**e**) in myeloid cell types (N=5 healthy donor PBMC) and CD4 and CD8+ T cells derived from patient PBMCs (n = 31). **g)** Representative images of T-cell lymphomas showing CD94 expression by immunohistochemistry (IHC). **h)** H-score for CD94 expression in various T-cell lymphoma biopsy samples from patients. mDC1, myeloid dendritic cells 1; mDC2, myeloid dendritic cell 2; pDC, plasmacytoid dendritic cell; NKTCL, NK/T-cell lymphoma; EATCL, enteropathy T-cell lymphoma; HSTCL, hepatosplenic T-cell lymphoma; TCL, T-cell lymphoma; CTCL, cutaneous T-cell lymphoma; PCGDTCL, primary cutaneous $\gamma\delta$ T-cell lymphoma; ATLL, adult T-cell leukemia/lymphoma.

**Extended Data Figure 2. Anti CD94 antibody generation. a)** L cells transduced with a lentiviral vector encoding human CD94 were characterized by flow cytometry. The plots show human CD94 expression on CD94-transduced L cells (right panel), compared to untransduced L cells (left panel). **b)** CD94-expressing L cells were injected into BALB/c mice at a dose of $5 \times 10^6$ cells via the footpad, twice a week for up to six weeks. Lymph nodes from both legs of immunized mice were collected for B cell isolation and fusion with Sp2/0 myeloma cells. Hybridoma cells were cultured in HAT medium for selection. Single clones were isolated by limiting dilution and characterized by ELISA and flow cytometry.

**Extended Data Figure 3. Cytotoxic activity and cytokine expression by CD94-targeting CAR T cells *in vitro*. a-b)** Cytotoxic activity by CD94-targeting CAR T cells (OX40 or CD28 co-stimulatory domain) against KHYG-1 clones with varying CD94 antigen density or CD94 knock-out at an Effector: Target (E:T) ratio of 1:1 was assessed by serial imaging using the Incucyte Live-Cell Analysis System over two days. **c)** Cytokine expression by CD94-targeting CAR T cells after 48 hours in response to KHYG-1 wild type (WT) or $CD94^{KO}$ KHYG-1 cells is shown as heatmap. Tumor alone and UTD T cells are shown as controls where applicable. Statistical significance was determined using one-way ANOVA with Dunnett's multiple comparisons test. ****: $p < 0.0001$. All experiments were performed in triplicate wells and the data represent mean ± standard error of mean where applicable. Results shown are representative of at least five separate experiments using CAR T cells generated from five different donors.

**Extended Data Figure 4. *In vivo* efficacy of CD94 CAR T cells against KHYG-1 tumor clone with low antigen density.** NSG mice were intravenously injected via the tail vein with $1 \times 10^6$ KHYG-1 clone 4 cells per mouse. After seven days, mice were untreated or treated with untransduced (UTD) T cells or CAR T cells at a dose of $4.5 \times 10^6$ CAR-T cells per mouse. **a-c)** Bioluminescence imaging (**a**), total flux (**b**), and body weight (**c**) are shown. Total flux values were calculated from normalized radiance spectra (p/s/cm²/sr) using Living Image software. Data are represented individually for each mouse. Statistical significance was determined using one-way ANOVA with Dunnett's multiple comparisons test (**: $p < 0.005$). UTD, untransduced; p/s, photons/second; ns, not significant.

**Extended Data Figure 5. Antitumor efficacy of CD94 CAR T cells against hepatosplenic T-cell lymphoma (HSTCL) patient-derived xenograft (PDX) model. a)** Experimental outline to assess efficacy of CD94 CAR T cells against PDX model of hepatosplenic T cell lymphoma (HSTCL) in NSG mice. **b** and **c)** Survival of mice (**b**) and body weight (**c**) in HSTCL PDX model. **d)** Spleens of CAR-T-treated and control groups at necropsy. **e)** Flow cytometry analysis of mouse tissue samples from CD94 CAR-T-treated and control groups to determine presence of tumor cells and human T cells. Statistical analysis for survival was determined using log-rank (Mantel-Cox) test (**: $p < 0.005$). Statistical significance for body weight was determined using one-way ANOVA with multiple comparisons test. Results shown are representative of at least three separate experiments using CAR T cells generated from three different donors. UTD, untransduced; ns, not significant.

**Extended Data Figure 6. Survival analysis of mice treated with rapidly manufactured CD94-targeting CAR T cells. a)** Schematic of experimental set-up to assess the effect of CD94 CAR T cells against NKL CDX model. Survival of mice treated with Day 3 CAR-T (**b**) vs. Day 10 CAR-T (**c**) CAR-T along with control groups. Statistical significance was determined using the log-rank (Mantel-Cox) test (**: $p < 0.005$).

**Extended Data Figure 7. Effects of CD94-targeting CAR T cells on immune cells *in vivo*. a-c)** Humanized NSG-IL15 mice engrafted with $CD34^+$ hematopoietic cord blood stem cells were untreated or treated with CD94 CAR T cells. Flow cytometry was performed to assess CD94 CAR-

T cell-mediated cytotoxicity against NK cells (**a**), CD94$^+$ T cells (**b**), and CD94$^+$ NK cells (**c**). **d-g)** Graphs show the proportional changes in total T and NK cells (**d-e**), and CD94$^+$ T cells (**f**) and CD94$^+$ NK cells (**g**), over the course of one week following CD94 CAR T cell treatment. Data are presented individually for each mouse. Statistical significance was determined using Student's *t*-test. ***: P < 0.0005; *: P < 0.05; ns: not significant.

**Extended Data Figure 8. Efficacy of dtBCMA safety switch efficacy *in vitro*. a-b)** CD94 CAR T cells harboring a dtBCMA safety switch construct were untreated or treated at the indicated concentrations with belantamab mafodotin (**a**) or elranatamab (**b**) or their respective isotype IgG antibody controls. CAR T cells at each indicated time point were determined by flow cytometry and representative plots from triplicate wells is shown. Results shown are representative of at least two separate experiments using CAR T cells generated from two different donors.

**Extended Data Figure 9. Specificity of dtBCMA safety switch and efficacy belantamab mafodotin *in vivo*. a)** CD94 CAR T cells without a dtBCMA safety switch were untreated or treated at the indicated concentrations with teclistamab. CAR T cells at each indicated time point were determined by flow cytometry and representative plots from triplicate wells is shown. **b)** NSG mice injected with CD94 CAR T cells with dtBCMA safety switch were treated with belantamab mafodotin or isotype control antibody on day 0 and representative dot plots showing CAR T cell percentage as determined by flow cytometry serially at the indicated time points in peripheral blood or in bone marrow and spleen at necropsy are shown. Plots are representative of one three mice from two independent biological experiments using CAR T cells derived from two different healthy donors.

Extended Data Figure 1

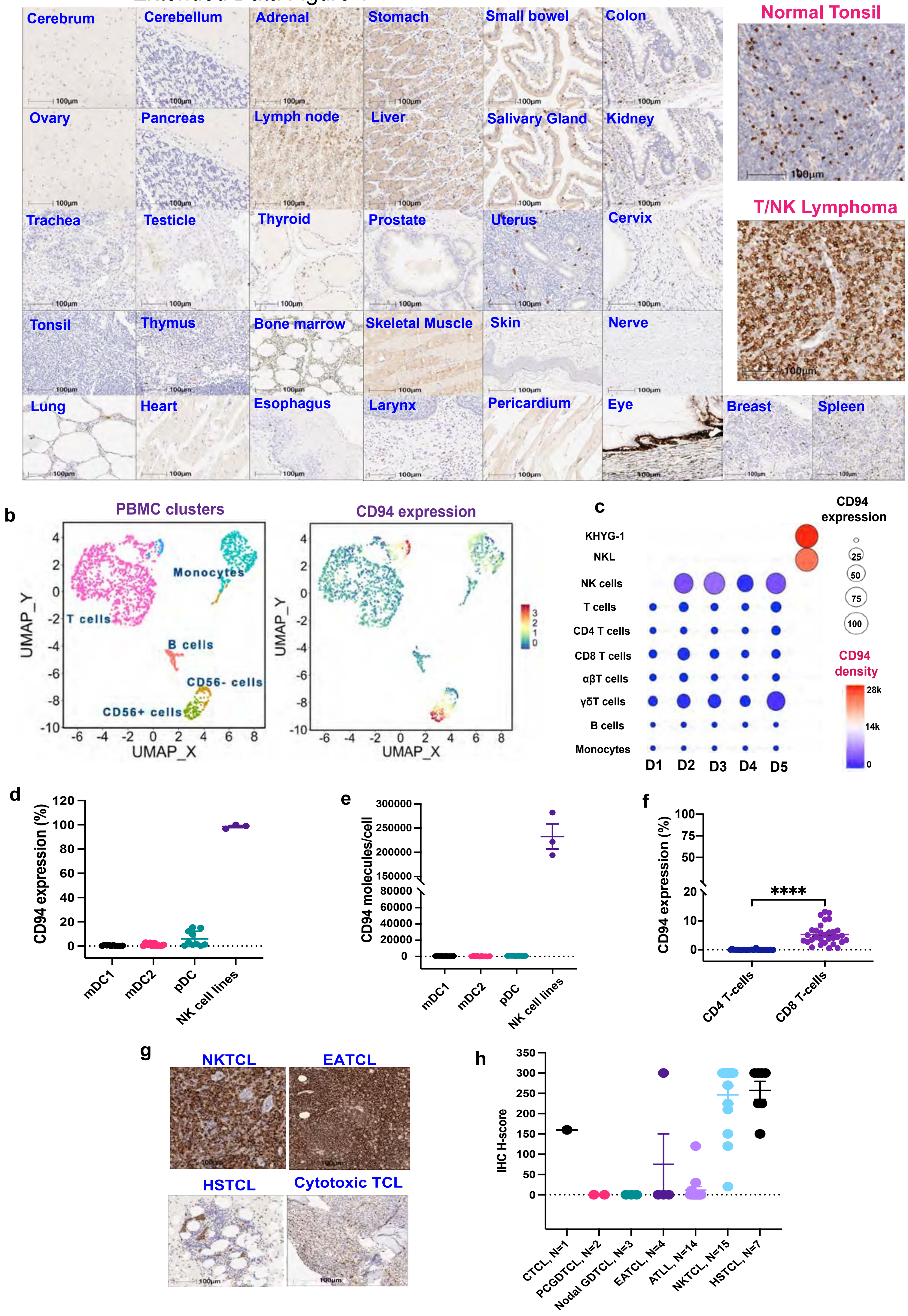

Extended Data Figure 2

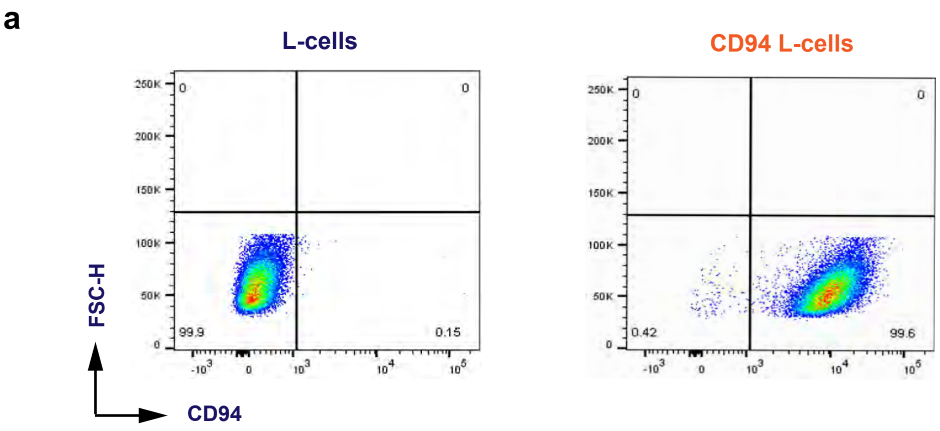


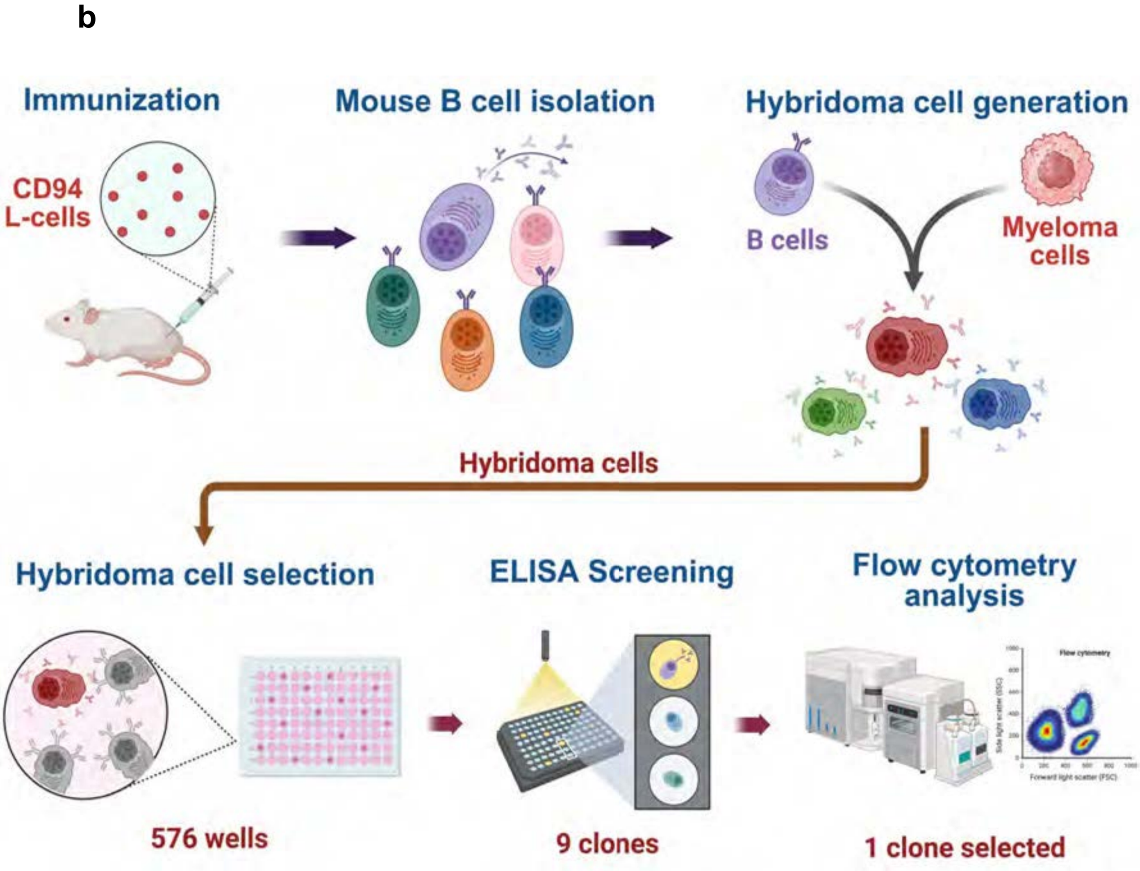

### Extended Data Figure 3

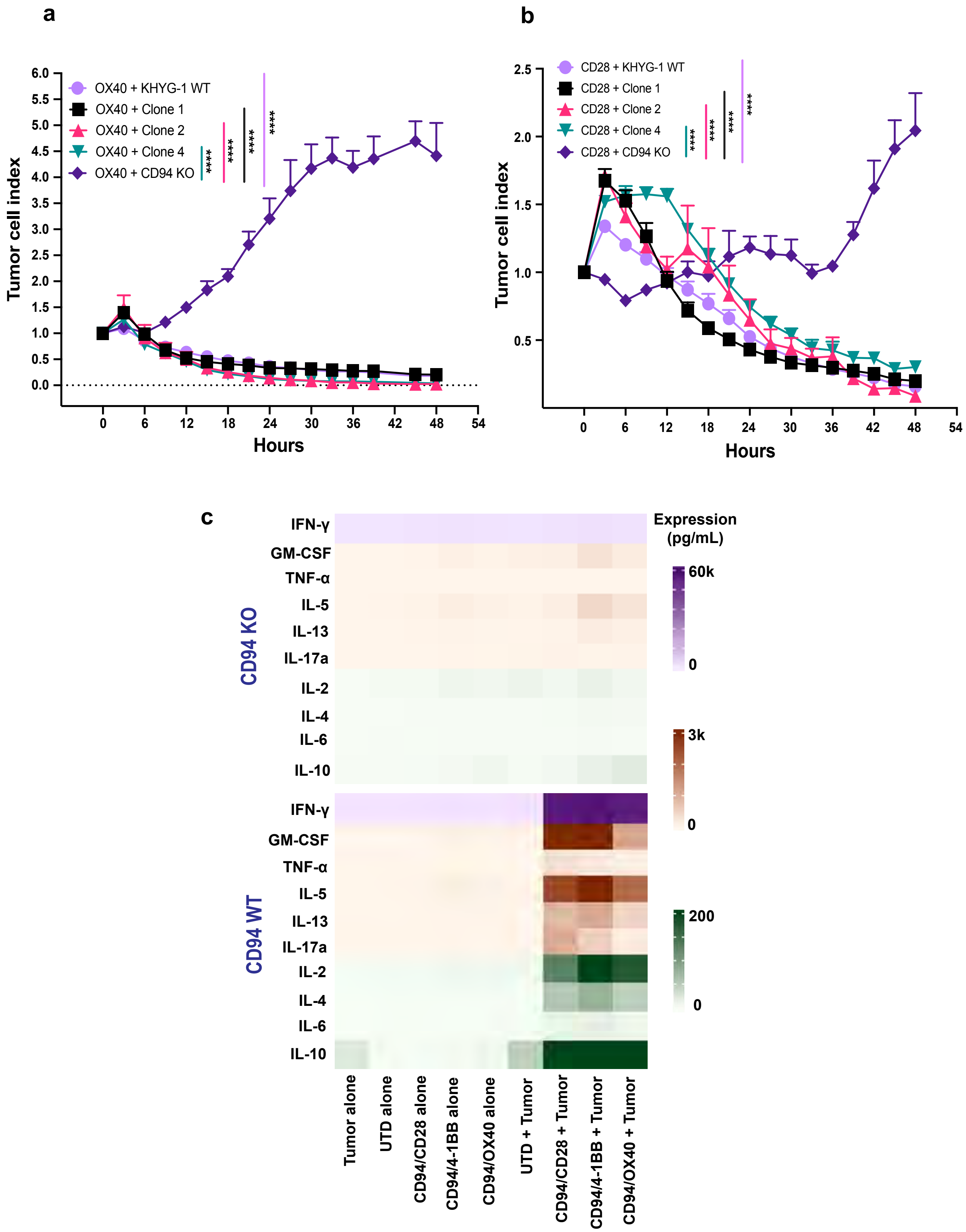

### Extended Data Figure 4

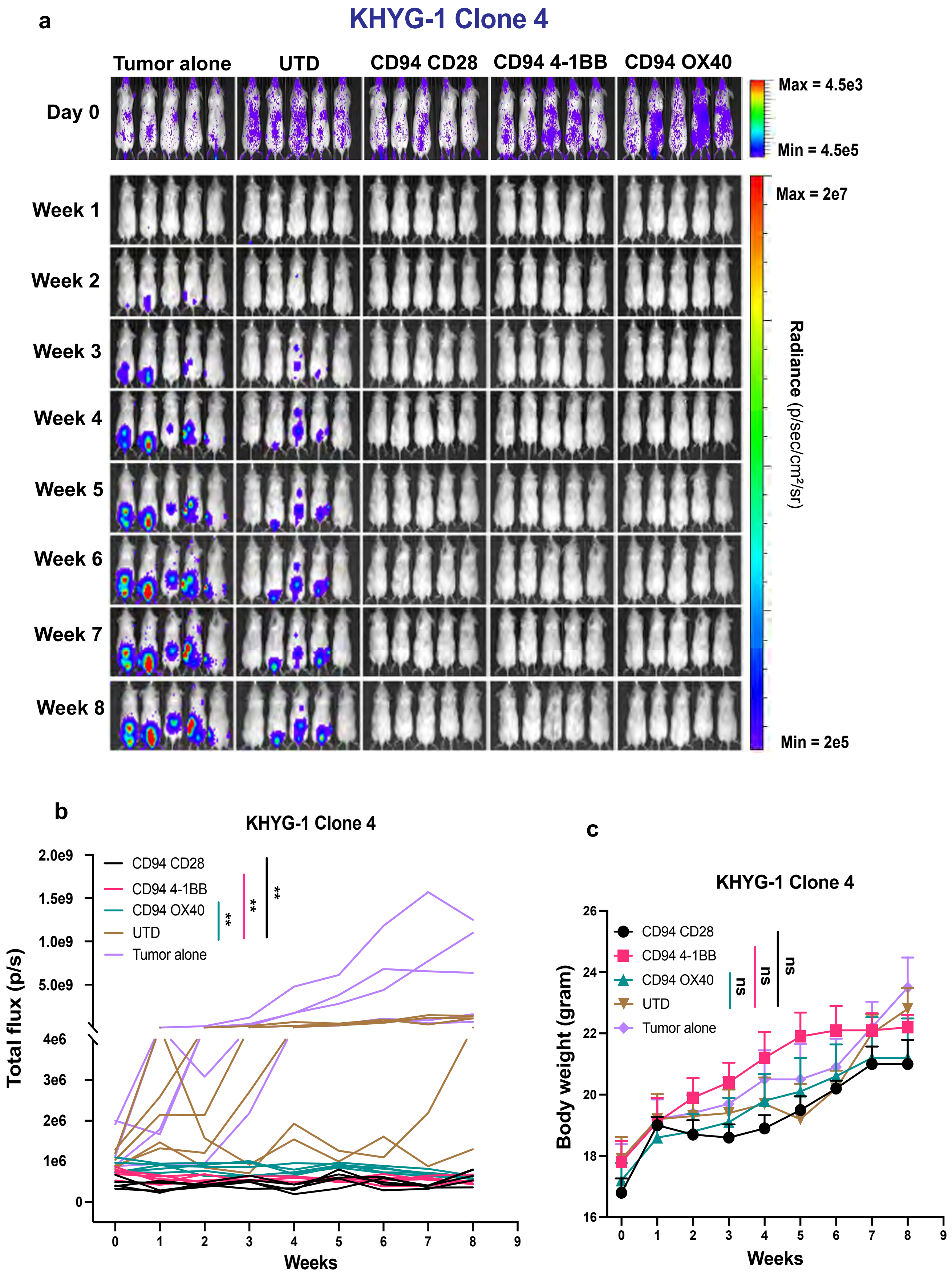

### Extended Data Figure 5

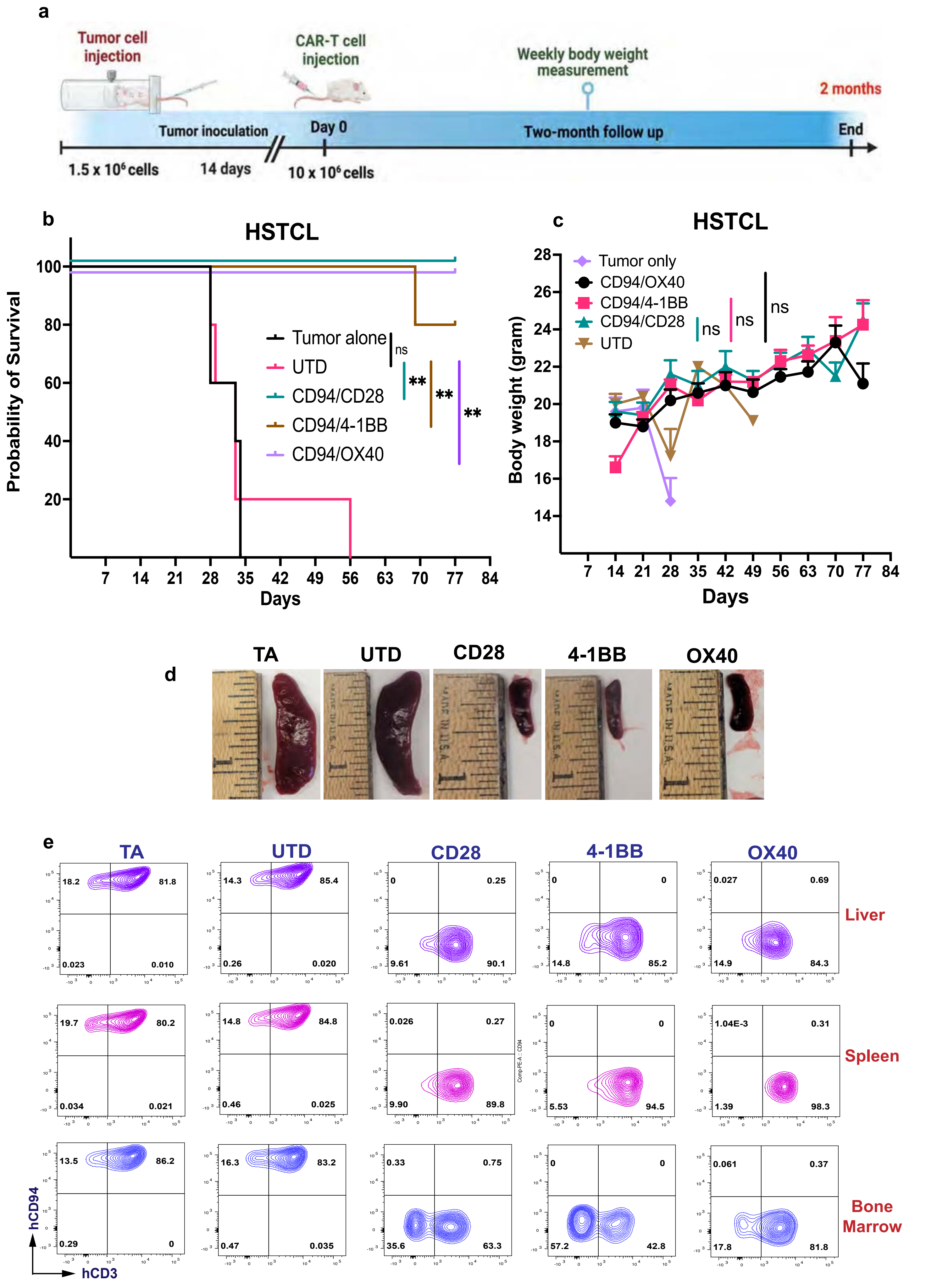

### Extended Data Figure 6

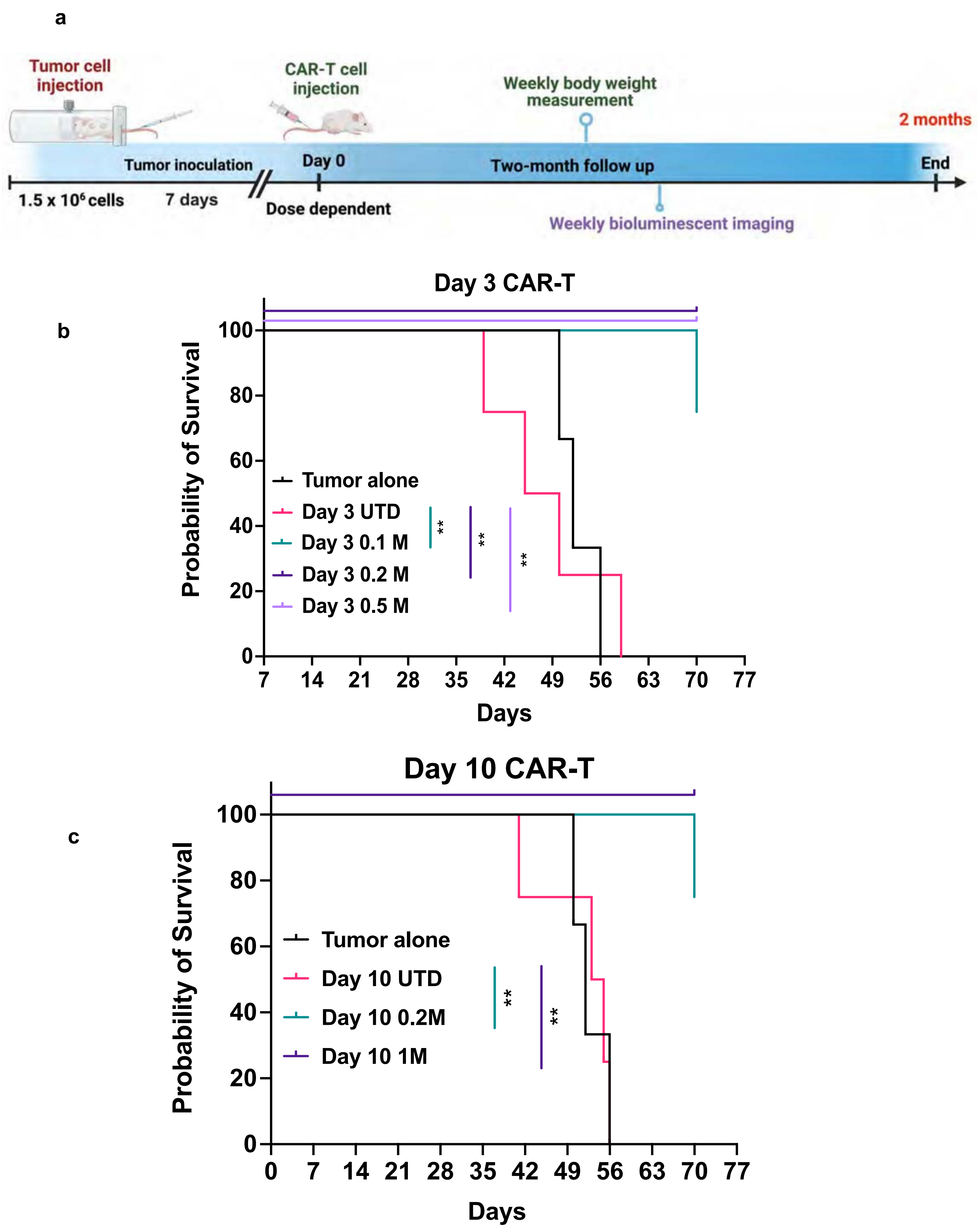

### Extended Data Figure 7

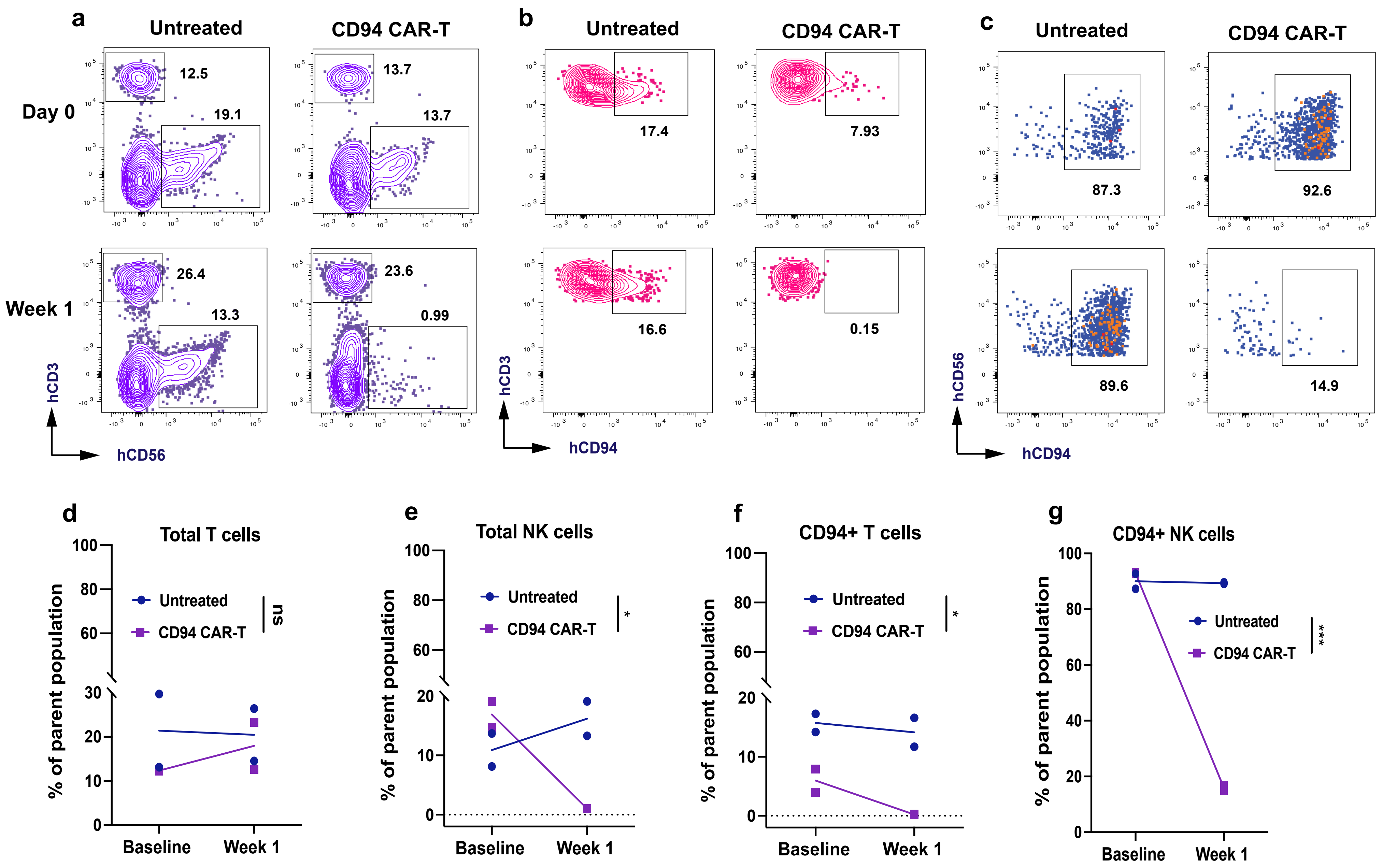

### Extended Data Figure 8

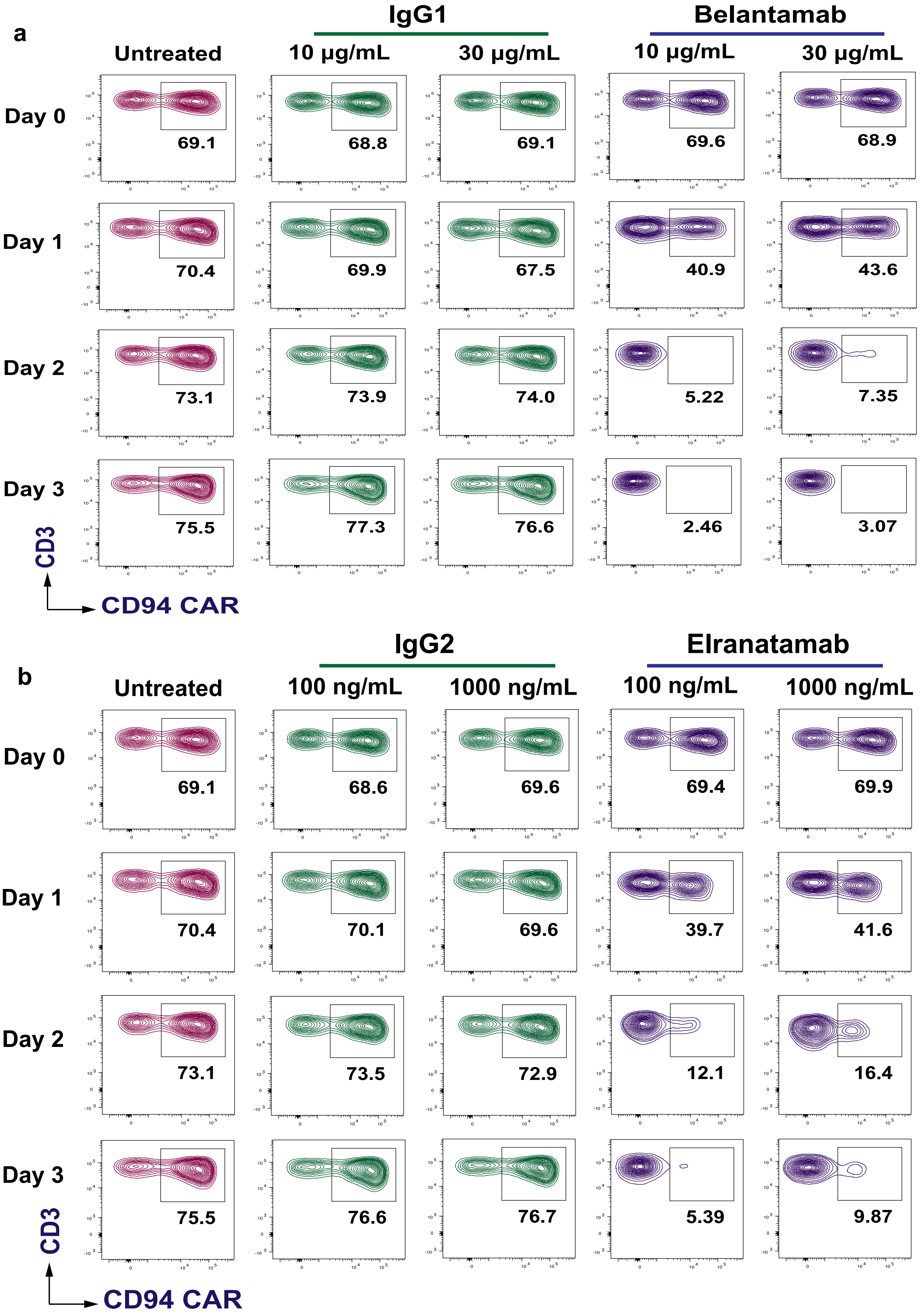

Extended Data Figure 9

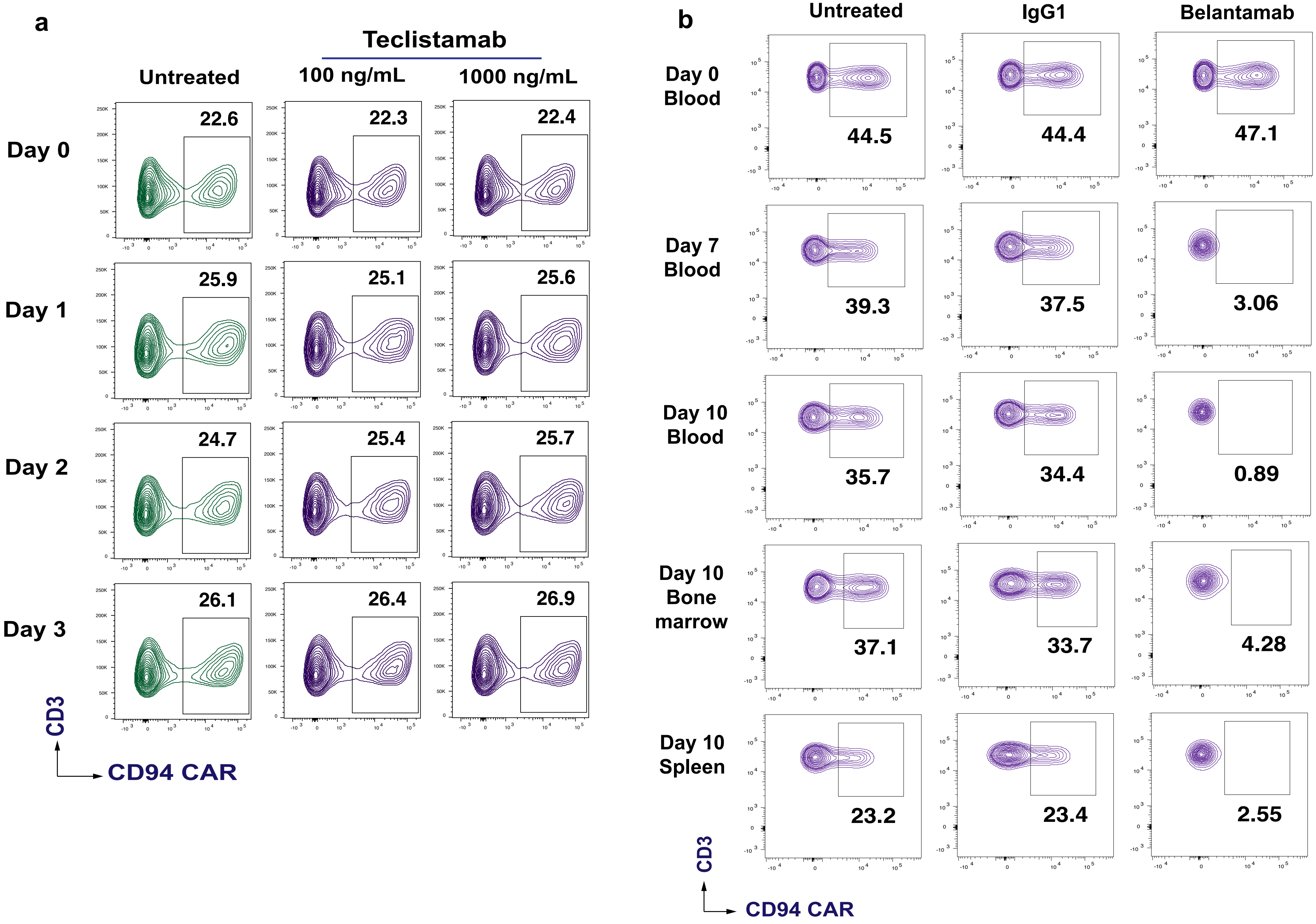
